# The genomics and evolution of inter-sexual mimicry and female-limited polymorphisms in damselflies

**DOI:** 10.1101/2023.03.27.532508

**Authors:** B. Willink, K. Tunström, S. Nilén, R. Chikhi, T. Lemane, M. Takahashi, Y. Takahashi, E. I. Svensson, C. W. Wheat

## Abstract

Sex-limited morphs can provide profound insights into the evolution and genomic architecture of complex phenotypes. Inter-sexual mimicry is one particular type of sex-limited polymorphism in which a novel morph resembles the opposite sex. While inter-sexual mimics are known in both sexes and a diverse range of animals, their evolutionary origin is poorly understood. Here, we investigated the genomic basis of female-limited morphs and male mimicry in the Common Bluetail damselfly. Differential gene expression between morphs has been documented in damselflies, but no causal locus has been previously identified. We found that male-mimicry originated in an ancestrally sexually-dimorphic lineage in association with multiple structural changes, probably driven by transposable element activity. These changes resulted in ∼900 kb of novel genomic content that is partly shared by male mimics in a close relative, indicating that male mimicry is a trans-species polymorphism. More recently, a third morph originated following the translocation of part of the male-mimicry sequence into a genomic position ∼3.5 mb apart. We provide evidence of balancing selection maintaining male-mimicry, in line with previous field population studies. Our results underscore how structural variants affecting a handful of potentially regulatory genes and morph-specific genes, can give rise to novel and complex phenotypic polymorphisms.

## Main

Sexual dimorphism is one of the most fascinating forms of intra-specific phenotypic variation in animals. Sexes often differ in size and colour, as well as the presence of elaborated ornaments and weaponry. Theoretical and empirical studies over many decades have developed a detailed framework of sexual selection and sexual conflict, explaining why these differences arise and how they become encoded in sex differentiation systems^1–3^. However, a growing number of examples of inter-sexual mimicry^4–7^ suggest that sexual dimorphism can be evolutionarily fragile and quite dynamic. Inter-sexual mimicry has evolved in several lineages, when individuals of one sex gain a fitness advantage, usually frequency– or density-dependent, due to their resemblance to the opposite sex. For example, males who mimic females, as seen in the Ruff (*Calidris pugnax*) and the Melanzona Guppy (*Poecilia parae*), forgo courtship and ‘sneak’ copulations from dominant males^4,5^, while females who mimic males, in damselflies and hummingbirds, avoid excessive male-mating harassment^6,8^. Inter-sexual mimicry thus requires the evolution of a novel sex-mimicking morph in a sexually-dimorphic ancestor. The occurrence of inter-sexual mimicry may be a intermediate step in the evolution of sexual monomorphism, it may be an ephemeral state, or it may be maintained as a stable polymorphism. In any case, sexual mimics harbour genetic changes that attenuate or prevent the development of sex-specific phenotypes, and can therefore provide insights into the essential building blocks of sexual dimorphism^9^.

Considerable research effort has been devoted to uncover the genetic basis of discrete phenotypic polymorphisms, such as those associated with alternative reproductive or life-history strategies^10–14^. Together, these studies highlight a vast diversity of mechanisms used by evolution to package complex phenotypic differences into a single locus that is protected from the eroding effects of recombination. On one extreme, phenotypic morphs may evolve via massive insertions, deletions, or inversions that lock together dozens to hundreds of genes into supergenes^15–17^. On the other end, much smaller structural variants (SVs), confined to a few thousand base pairs, can modulate the expression of one or a few regulators of pleiotropic networks, resulting in markedly different morphs^11,12,18^. We are clearly only starting to get a glimpse of the major themes among these genetic mechanisms. For example, it is not known whether genomic architecture determines the type and breadth of co-varying traits or the likelihood of polymorphisms evolving in specific lineages^19^.

A few of these studies have focused on sex-limited polymorphisms, where one of the morphs shares the overall appearance, such as the colour pattern, of the opposite sex^10,14,20^. Such sex-limited morphs may illustrate novel origins of sexual dimorphism, driven by either sexual selection in males^14^ or natural selection in females^18,21^. Alternatively, sex-limited polymorphisms may arise with the evolution of inter-sexual mimicry. Crucially, empirical support for the evolution of inter-sexual mimicry demands both a macroevolutionary context for the polymorphism, showing that sexually dimorphism is ancestral, and a documented advantage of sexual mimics in at least some social contexts. There is therefore a need to integrate genomic, microevolutionary and phylogenetic evidence into our understanding of the evolutionary dynamics of sexual dimorphism and inter-sexual mimicry. This integrative approach has been overall rare, and applied mostly to the study of alternative male reproductive strategies^18,22^. Yet, female mimicry of males may be more common than historically appreciated^23^, and the genetic basis of such mimicry remains largely unexplored^24–26^.

The Common Bluetail damselfly *Ischnura elegans* (Odonata) has three female-limited morphs (namely *O*, *A* and *I*) that differ in colouration, whereas males are always monomorphic^27^. *O* females display the colour pattern and developmental colour changes inferred as ancestral in a comparative analysis of the genus *Ischnura*^28^ (Fig. 1). Male-like (*A*) females are considered male mimics, who experience a frequency-dependent advantage of reduced male-mating and premating harassment due to their resemblance to males^6^. Finally, the *I* morph shares its stripe pattern and immature colouration with the *A* morph^27^ (Fig. 1), but develops a yellow-brown background colouration with age, eventually resembling the *O* morph upon sexual maturation^29^. *I* females are only known in *I. elegans* and a few close relatives^28^ (Fig. 1), and their evolutionary relationship to *A* and *O* females remains unresolved. The behaviour, ecology, and population biology of *I. elegans* have been intensely investigated for over two decades, making it one of the best understood female-limited polymorphisms, in terms of how morphs differ in fitness-related traits and how alternative morphs are maintained sympatrically over long periods^30–33^.

**Figure 1.**
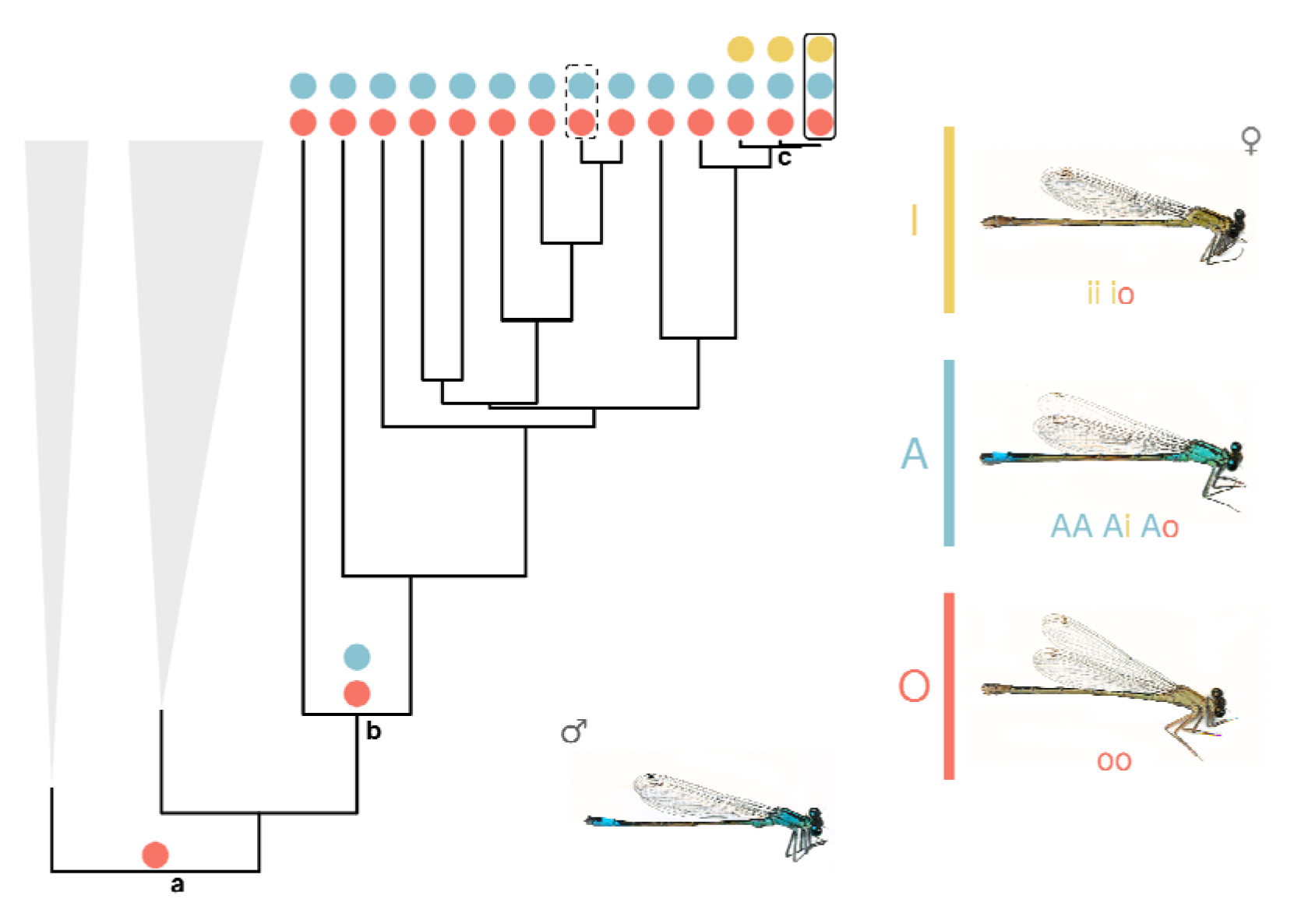
The evolution of female-limited colour polymorphisms in *Ischnura* damselflies. **a** A previous phylogenetic study and ancestral state reconstruction^28^ proposed that the genus *Ischnura* had a sexually dimorphic ancestor, with *O*-like females (red circle). The *O* morph is markedly different from males, having a bronze-brown thorax and faint stripes, instead of the black thoracic stripes on a bright blue background of males. **b** Male mimicry (*A* females, blue circle) has evolved more than once, for instance, in an ancestor of the (expanded) clade that includes the Common Bluetail (*I. elegans*, encircled with solid line) and the Tropical Bluetail (*I. senegalensis*, encircled with dashed line). **c** *I. elegans* is a trimorphic species, due to the recent evolution of a third female morph, *I* (yellow circle). In *I. elegans*, morph inheritance follows a dominance hierarchy, where the most dominant allele produces the *A* morph and two copies of most recessive allele are required for the development of *O* females. In contrast, the *O* allele is dominant in *I. senegalensis*^106^. Terminal nodes in the phylogeny represent different species. Gray triangles represent other clades of *Ischnura* that are collapsed for clarity.

Nonetheless, the molecular basis of this polymorphism remains unknown.

To advance our understanding of the evolution of complex phenotypes, such as sexual dimorphism and sex-specific morphs, we identify the genomic region responsible for the female-limited colour polymorphism in *I. elegans*. Using a combination of reference-based and reference-free genome wide association studies (GWAS), upon morph-specific genome assemblies, we revealed two novel regions adding up to ∼900 kb, that are associated with the evolutionary origin of the male-mimicking *A* morph. These structural variants, probably generated and expanded by transposable element (TE) activity, are partly shared by male-mimicking females of the Tropical Bluetail damselfly (*Ischnura senegalensis*), indicating that male mimicry is a trans-species polymorphism. We also show that the novel *I* morph evolved via an ectopic recombination event, where part of the *A*-unique genomic content was translocated into an *O* genomic background. Finally, we examined the evolutionary dynamics of the colour morph locus and explored expression patterns of genes located in this region. Together, our results indicate that structural variation affecting a handful of genes and maintained by balancing selection provides the raw material for the evolution of a male-mimicking phenotype in pond damselflies.

## Results

### Male mimicry is encoded by a locus with a signature of balancing selection

We started by conducting three reference-based GWAS, comparing all morphs against each other in a pairwise fashion (Extended Data Fig. 1). We used an *A* morph genome assembly (Supporting Text 1) as mapping reference because structural variant analyses revealed that *A* females harbour genomic content that is absent in the other two morphs (see *Female morphs differ in genomic content* below). The draft assembly was scaffolded against the Darwin Tree of Life (DToL) reference genome to place the contigs in a chromosome level framework^34^. The DToL reference genome contains the *O* allele (see Supporting Text 2) and is assembled with chromosome resolution, except for chromosome 13, which is fragmented and consists of one main and several unlocalized scaffolds.

All pairwise GWAS between morphs pointed to one and the same unlocalized scaffold of chromosome 13 as the causal morph locus (Fig. 2a). Closer examination of this scaffold revealed two windows of elevated divergence between morphs (Fig. 2b). First, a narrow region near the start of the scaffold (∼50 kb – 0.2 mb) captures highly significant SNPs in both *A* vs *O* and *I* vs *O* comparisons (Fig. 2b). Thereafter and up to ∼1.5 mb, an abundance of SNPs differentiates *A* females from both *O* and *I* females, especially between ∼0.6 and ∼1.0 mb (Fig. 2b). These results are mirrored by Fst values across both regions (Fig. 2c).

**Figure 2.**
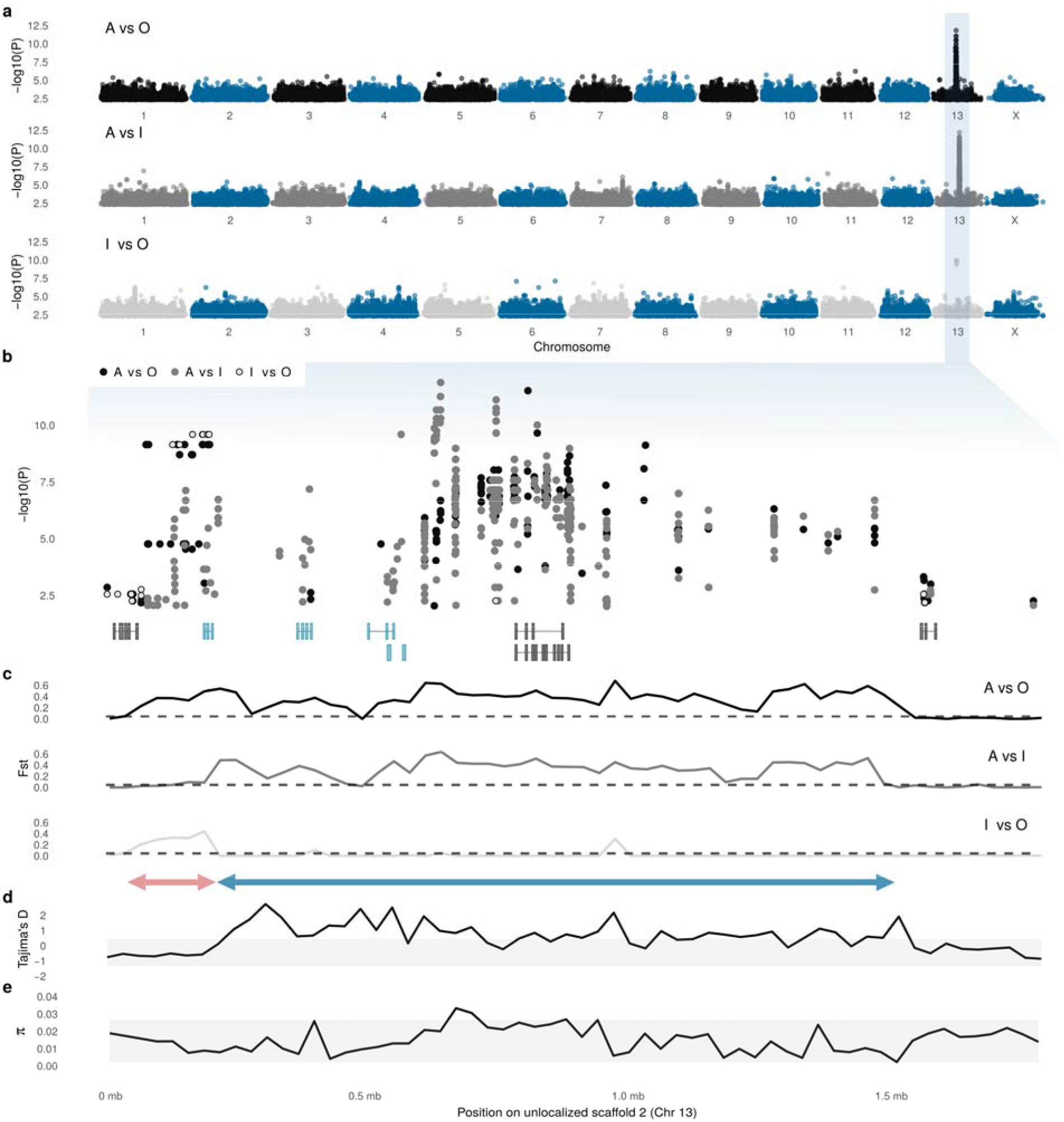
Morph determination in the Common Bluetail Damselfly (*Ischnura elegans*) is controlled in a ∼ 1.5 mb region of chromosome 13. **a** SNP-based genome-wide associations in all pairwise analyses between morphs. Genomic DNA from 19 wild-caught females of each colour morph and of unknown genotype was extracted and sequenced for these analyses. Illumina short reads were aligned against an *A* morph genome assembly, generated from Nanopore long-read data (Extended Data Fig. 1). **b** A closer look of the SNP associations on the unlocalized scaffold 2 of chromosome 13, which contained all highly significant SNPs. Transcripts expressed in at least one adult of both *I. elegans* and *I. senegalensis* are shown at the bottom (see also Fig. 6). Grey transcripts are shared by all morphs, whereas blue transcripts are uniquely present in *A* or *A* and *I* samples (see *Shared and morph-specific genes reside in the morph locus*). The *y* axis in **a** and **b** indicates unadjusted –Log_10_ P-values calculated from chi-squared tests. **c** Fst values averaged across 30 kb windows for the same pairwise comparisons as in the SNP based GWAS. The dashed line marks the 95 percentile of all non-zero Fst values across the entire genome. The red double arrow shows the region of elevated divergence between *O* and both *A* and *I* samples (∼50 kb – 0.2 mb). The blue double arrow shows the region of elevated divergence between *A* and both *O* and *I* samples (∼0.2 mb – 1.5 mb). Population-level estimates of **d** Tajima’s D, and **e** nucleotide diversity (π) averaged across 30 kb windows. The shaded area contains the 5-95 percentile of all genome-wide estimates.

Next, we investigated whether the morph locus carries a signature of balancing selection, as suggested by previous field studies of morph-frequency dynamics^31^. The larger genomic window that uniquely distinguishes *A* females from both *I* and *O* females displays a signature of balancing selection, indicated by highly positive values of Tajima’s D, exceeding the 95 percentile of genome-wide estimates (Fig. 2d). Conversely, values of both Tajimas’s D and π in the narrower window that differentiates *O* females from both *A* and *I* females (∼50 kb – 0.2 mb) fall within the 95 percentile of genome-wide estimates (Fig. 2d-e).

### Female morphs differ in genomic content

Previous studies have found that complex phenotypic polymorphisms are often underpinned by structural variants (SVs), arising from genomic rearrangements such as insertions, deletions and inversions^10,13,15,20^. As these variants can be difficult to detect in a reference-based analysis, we employed a *k*-mer based GWAS approach^35^ (Extended Data Fig. 1), which enables reference-free identification of genomic divergence between morphs. Significant *k*-mers in these analyses could represent regions that are present in one morph and absent in the other (i.e. insertions or deletions), or regions that are highly divergent in their sequence (as in a traditional GWAS).

First, we investigated the divergence associated with the male-mimicking *A* morph. Pairwise analyses revealed 568,039 and 508,031 *k*-mers (length = 31 bp) significantly associated with the *A* vs *O* and *A* vs *I* comparisons, respectively. To determine whether the associated *k*-mers represent differences in genomic content or sequence between the morphs, we mapped these *k*-mers to morph-specific reference genomes. If the associated *k*-mers are due to novel sequences found in one morph but not the other, we would expect a vast majority of the significant *k*-mers to be found in only one of the two morphs in a pairwise comparison. If the significant *k*-mers are instead owed to point mutations in high-identity sequences, there should be morph-specific *k*-mers in both morphs.

Most (> 98%) of the mapped *k*-mers in the *A* vs *O* and *A* vs *I* comparisons, aligned perfectly to a single ∼1.5 mb region of the unlocalized scaffold 2 of chromosome 13, in the *A*-morph assembly (Fig. 3a; Extended Data Table 1). This is the same region of the *A*-morph assembly that was previously identified in the standard GWAS (Fig. 2). In contrast, only ∼0.3% of the associated *k*-mers in the *A* vs *O* comparison were found anywhere in the *O*-assembly, and similarly, only ∼0.2% of the significant *k*-mers in the *A* vs *I* analysis mapped to the *I* assembly (Extended Data Table 1). These results thus suggested that a large region of genomic content is unique to the *A* haplotype.

**Figure 3.**
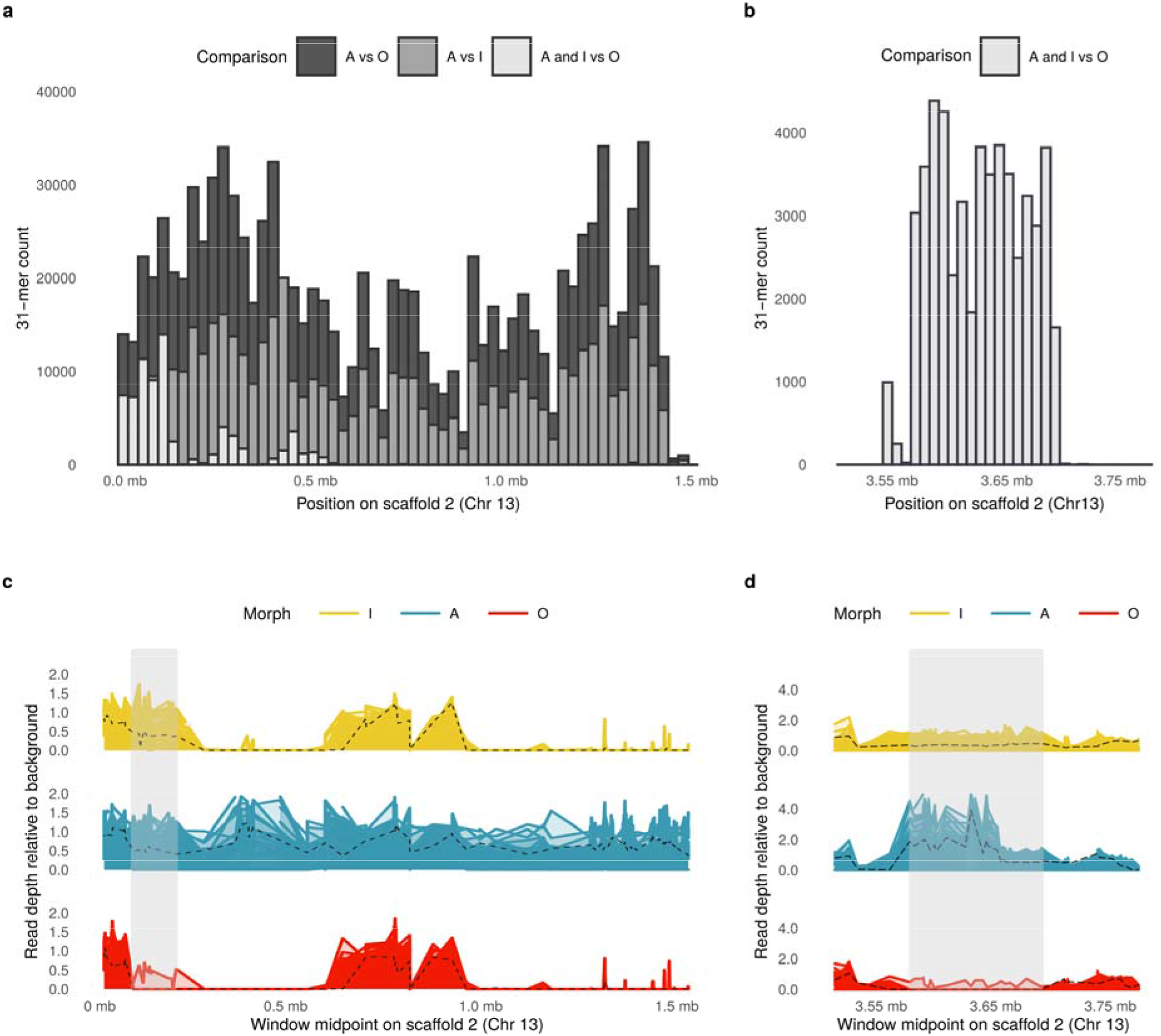
Female morphs of *Ischnura elegans* differ in genomic content. Number of significant *k*-mers (below the 5% false-positive threshold, see Methods) associated with pairwise genome-wide analyses and mapped to the unlocalized scaffold 2 of chromosome 13, in **a** the *A* morph assembly, and in **b** the *I* morph assembly. Standardized read depths along the unlocalized scaffold 2 of chromosome 13, relative to background coverage of **c** the *A* morph assembly, and **d** the *I* morph assembly. Solid lines (yellow, blue and red) show short-read data (19 samples per morph) and black dashed lines represent long-read data (1 sample per morph).Grey areas show regions of genomic content present in *A* and *I* individuals, but absent in all but one *O* sample. Note that different regions of the scaffold are plotted for the two assemblies (see main text).

Given that *A* and *I* females share their immature colour pattern^29,36^, we then tested for *k*-mer associations that would distinguish both *A* and *I* females from *O* females and found 85,134 such *k*-mers (Extended Data Table 1). When mapped to the *A* assembly, a majority of these *k*-mers were found near the start of the unlocalized scaffold 2 of chromosome 13 (Fig. 3a), where we previously reported pronounced divergence of *O* females (Fig. 2b-c). However, when mapped to the *I* assembly, most of the significant *k*-mers were found in a different region of the same scaffold, separated by approximately 3.5 mb (Fig. 3b). These results thus suggested that *A* and *I* females share genomic content that is absent in *O*. However, in the *I* haplotype this content occupies a different chromosomal location.

To further investigate the distribution of genomic content among morphs, we plotted the standardized number of mapped reads (read depths) along the ∼1.5 mb region of the *A* assembly that included most of the significant *k*-mers (Extended Data Fig. 1). Here, we expected read depth values around 0.5 (heterozygous) or 1.0 (homozygous) for all *A* samples, whereas *I* and *O* samples should have read depths of 0, if genomic content is uniquely present in the *A* allele (because *I* and *O* individuals lack the *A* allele, Fig. 1). Read depths confirmed that male-mimicking *A* females are differentiated by genomic content. Specifically, there are two windows of the *A* assembly (of ∼400 kb and ∼500 kb) where no *I* or *O* data maps to the assembly after filtering repetitive sequences (Fig. 3c), and which are therefore uniquely present in *A* females.

These two windows of *A*-specific content are separated by a region between ∼0.6 and ∼1.0 mb that is shared among all morphs (Fig. 3c), and highly divergent in SNP-based comparisons involving the *A* morph (Fig. 2b). Finally, the region including most significant *k*-mers in the *A* and *I* vs. *O* comparison is present in all *A* and *I* samples but absent in all *O* samples, except for one individual (Fig. 3c; Supporting Text 3). As noted in the *k*-mer GWAS, this region of genomic content shared by *A* and *I* individuals is located in different regions, separated by ∼3.5 mb, in the two assemblies (Fig. 3d).

By combining reference-based GWAS, reference-free GWAS and read-depth approaches, we have identified three haplotypes controlling morph development in the Common Bluetail. The *A* and *I* haplotypes share ∼150 kb that are absent in *O*. The *A* haplotype has two additional windows of unique genomic content, adding up to ∼900 kb. In the *A* haplotype, a single ∼1.5 mb window (hereafter the morph locus) thus contains the regions of unique genomic content, the region exclusively shared between *A* and *I*, and the SNP-rich region present in all morphs. In the *I* haplotype the region exclusively shared with *A* occupies a single and different locus separated by about 3.5 mb (Fig. 4a). These large and compounded differences in genomic content between haplotypes suggest that multiple structural changes on a multi-million base-pair region were responsible for the evolution of novel female morphs in *Ischnura* damselflies.

**Figure 4.**
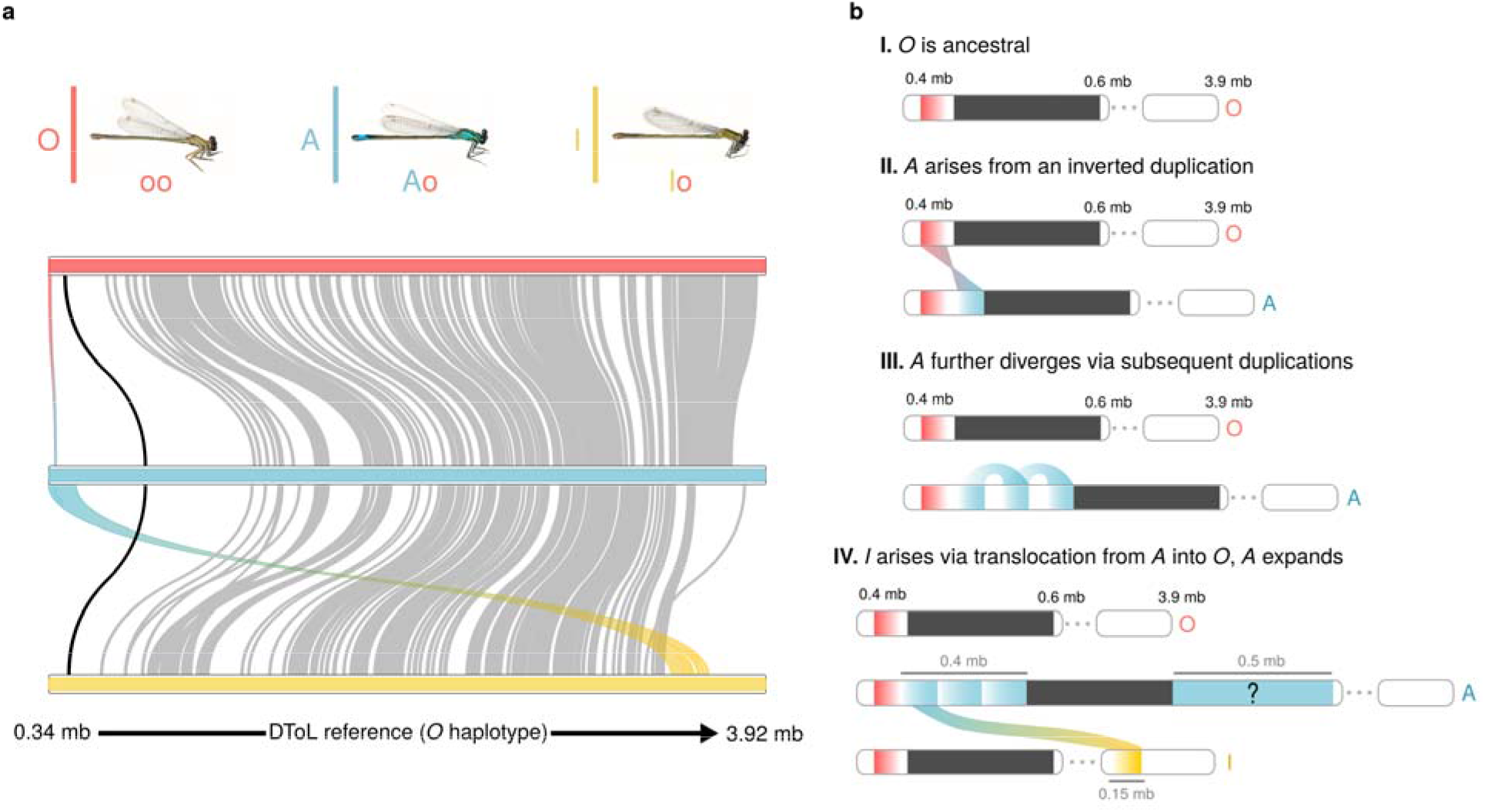
Structural variants differentiate morph haplotypes in the Common Bluetail Damselfly (*Ischnura elegans*). **a** Alignment between morph-specific genomes assembled from long-read Nanopore samples with genotypes *Ao*, *Io*, and *oo*. Grey lines represent alignments of at least 5 kb and > 70% identity. The black line connects regions of genomic content shared by the three morphs within the morph locus. The red to blue gradient represents a ∼20 kb region that carries an inversion signature in *A* and *I* females relative to the *O* haplotype (see Extended Data Fig. 2). The blue to yellow gradient represents a ∼ 150 kb alignment between the start of the unlocalized scaffold 2 of chromosome 13 in *A* and a region ∼ 3.5 mb apart in the *I* haplotype. Coordinates at the bottom are based on the Darwin Tree of Life (DToL) reference assembly. **b** Schematic illustration of the hypothetical sequence of events responsible for the evolution of novel female morphs. First, a sequence originally present in *O* was duplicated and inverted in tandem, potentially causing the initial divergence of the *A* allele. Second, part of this inversion was subsequently duplicated in *A*, in association with a putative TE, leading to multiple inversion signatures in the *A* haplotype relative to an *O* reference (see Extended Data Fig. 3). Finally, part of the *A* duplications were translocated into a position ∼ 3.5 mb downstream into an *O* background, giving rise to the *I* morph. Currently, *A* females are also characterized by another region of unique content and unknown origin (question mark). *A* female show elevated sequence divergence in the internal region of the morph locus that is shared by all haplotypes (dark grey bars, see also black line in **a**). Coordinates on the *O* haplotype are based on the (DToL) reference assembly. Grey numbers in IV give the approximate size of genomic sequences in *A* and *I* that are absent in *O*.

### TE propagation and recombination likely explain the origins of novel female morphs

Based on previous inferences of the historical order in which female morphs evolved (Fig. 1), we hypothesized that genomic divergence first occurred between *O* and *A* females, with some genomic content being then translocated from *A* into an *O* background, leading to the evolutionary origin of *I* females. We analyzed structural variants between morphs to test this hypothesis (Extended Data Fig. 1; Supporting Text 4), and uncovered evidence of a ∼20 kb sequence in the *O* haplotype that is duplicated and inverted in tandem in derived morphs (*A* and *I*; Fig. 4b; Extended Data Fig. 2). An investigation of the reads mapping to the inversion breakpoints suggested that additional duplications in the *A* genome, presumably via TE proliferation, may be related to the evolution of inter-sexual mimicry (Fig. 4b; Extended Data Fig. 3). Interestingly, TE content is enriched and recombination is reduced not just in the vicinity of the morph locus, but across the entire chromosome 13 (Extended Data Fig. 4-5; Supporting Text 4). Finally, evidence of a translocation of an *A*-derived genomic region back into an *O* background (Extended Data Fig. 6; Supporting Text 4) implied that the *I* morph evolved from an ectopic recombination event between between *A* and *O* morphs (Fig. 4b). This scenario is also consistent with our previous *k*-mer GWAS and read-depth results, where we found that the only region differentiating both *A* and *I* females from *O* females is located ∼3.5 mb in the *I* haplotype.

### Male mimicry is a trans-species polymorphism

Ancestral state reconstruction of female colour states had previously pointed to an ancient origin of male mimicry in the clade that includes *I. elegans* and several other widely-distributed *Ischnura* damselflies^28^ (Fig. 1). We investigated whether male mimicry is in fact a trans-species polymorphism using *de novo* genome assemblies from the closely related Tropical Bluetail (*Ischnura senegalensis*) (Extended Data Fig. 1). *I. senegalensis* shares a common ancestor with *I. elegans* about 5 Ma^28^, and has both a male-mimicking *A* morph and a non-mimicking morph, which resembles the *O* females of *I. elegans*^28,37^ (Fig. 5a).

**Figure 5.**
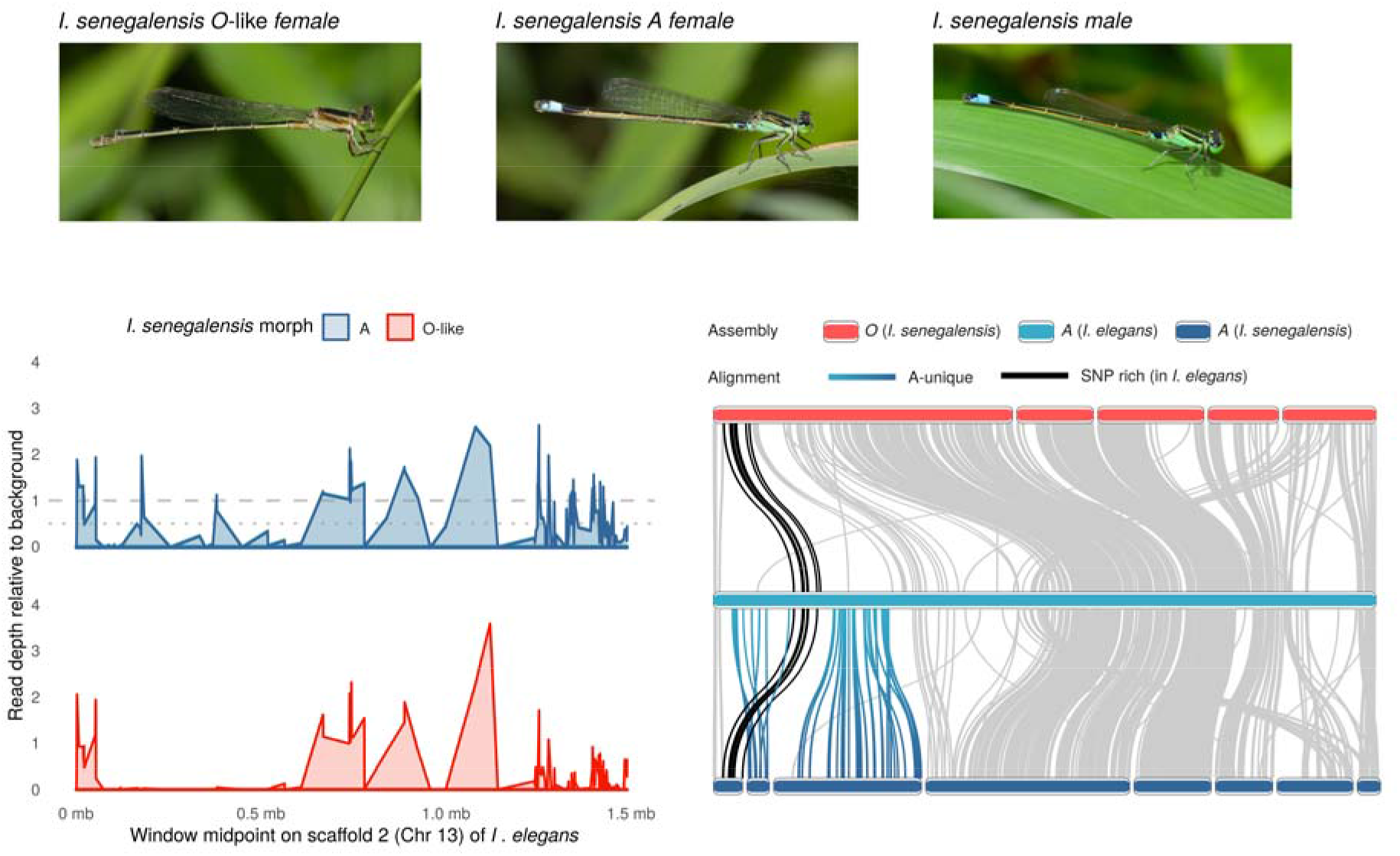
A shared genomic basis of *A* females in *Ischnura elegans* and *Ischnura senegalensis* **a** *I. senegalensis* is a female-dimorphic species, where one female morph (*O*-like) is distinctly different from males and resembles *O* females in *I. elegans*, and the other female morph (*A*) is a male-mimic. Photo credit: Mike Hooper. **b** Standardized read depth of pool-seq samples (n = 30 females of each morph per pool) of *I. senegalensis*, against the *A* morph assembly of *I. elegans*, calculated in 500 bp windows. The x-axis shows the first 1.5 mb of the unlocalized scaffold 2 of chromosome 13. **c** Alignments between morph-specific genomes from a homozygous *O*-like female of *I. senegalensis* (top), an *Ao* female of *I. elegans* (middle) and a homozygous *A* female of *I. senegalensis* (bottom). Lines connecting the assemblies represent alignments of at least 500 bp and > 70% identity. The black line connects genomic content in the morph locus, which is shared by the three morphs of *I. elegans*. In *I. elegans*, this region is rich in SNPs differentiating *A* females from the other two morphs (see Fig. 2b). The blue-turquoise gradient connects sequences uniquely present in the *A* morphs of *I. elegans* and *I. senegalensis*.

We reasoned that if morph divergence is ancestral, the genomic content that is uniquely present in *A* females or shared by *A* and *I* females in *I. elegans* should be at least partly present in *A* females of *I. senegalensis*, but absent in the alternative *O*-like female morph (see Supporting Text 5). This prediction was supported by differences in standardized read depths between the *A* and *O*-like pool of *I. senegalensis*, specifically at the morph locus of *I. elegans* (Fig. 5b; Supporting Text 5). A shared genomic basis of inter-sexual mimicry for the two species was also supported by the same ∼20 kb inversion signature in the *A* pool against an *O* assembly, as detected in *A* and *I* females of *I. elegans* (Extended Data Fig. 7). Finally, assembly alignments between *O*-like and *A* haplotypes of *I. senegalensis* showed that the *A*-specific genomic region of *I. elegans* is partly present in the *A* but not the *O*-like assembly of *I. senegalensis* (Fig. 5c).

### Shared and morph-specific genes reside in the morph locus

Finally, we examined gene content and expression patterns in the morph locus. As female morphs differ in genomic content as well as sequence, the phenotypic effects of the morph locus could come about in at least three non-exclusive ways. First, entire gene models may be present in some morphs and absent in others. Second, genes present in all morphs may differ in expression patterns. Third, genes may encode different amino acid sequences in different female morphs. We used newly generated and previously published^38^ RNAseq data to investigate these questions (Extended Data Fig. 1), and capitalized on the annotations of the reference genome of *I. elegans*^34^, as well as transcripts assembled *de novo* in our *A*-morph genome assembly. Because the genetic basis of inter-sexual mimicry is shared between *I. elegans* and *I. senegalensis* (Fig. 5), we focus on genes that are expressed in both species in at least one individual (Fig. 6a).

**Figure 6.**
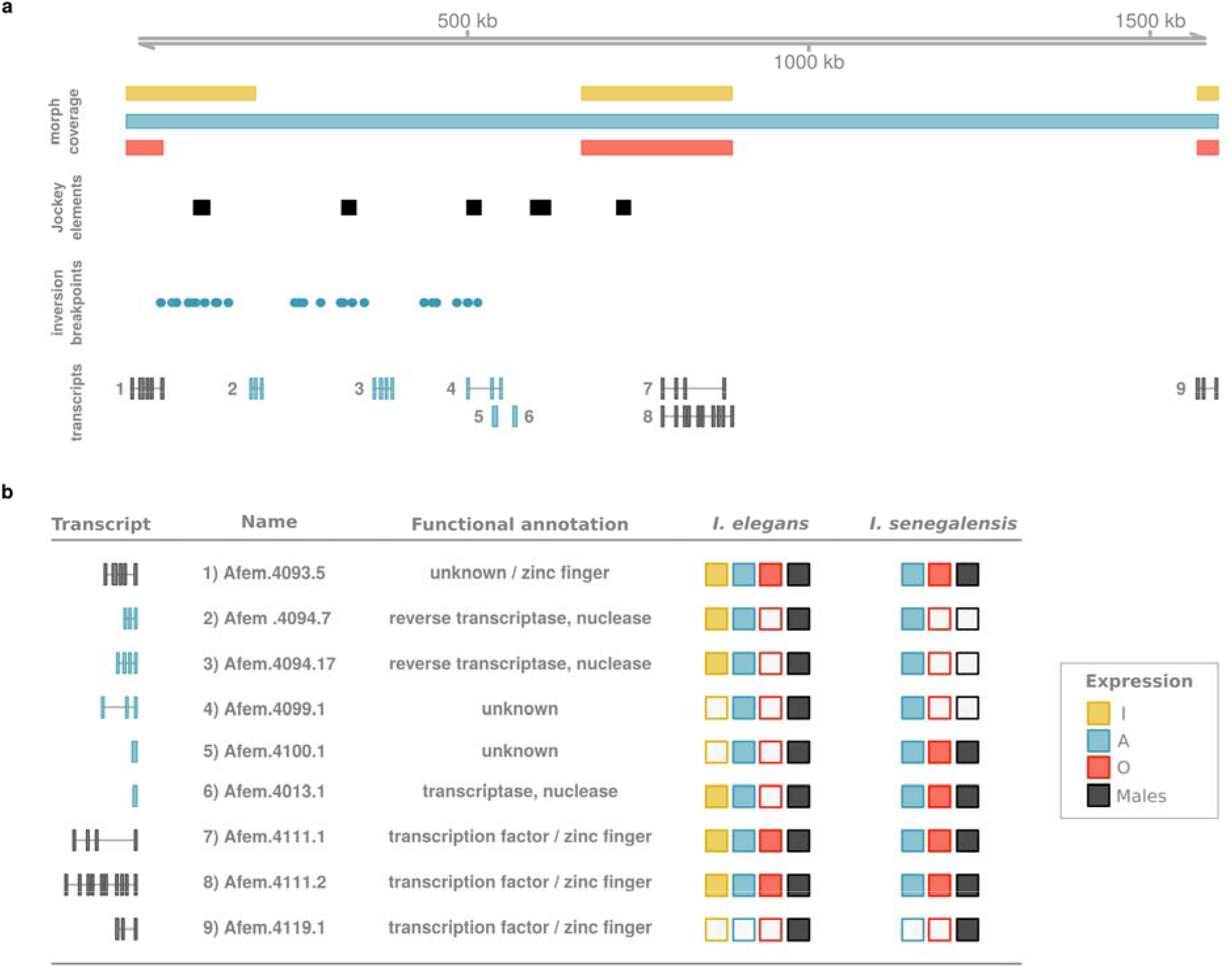
The morph locus of *Ischnura elegans* is situated in the unlocalized scaffold 2 of chromosome 13 **a** Diagram of the ∼ 1.5 mb morph locus on the *A*-morph assembly, showing from top to bottom: morph-specific read depth coverage, the location of LINE retrotransposons in the the Jockey family, the mapping locations of *A*-derived reads with a previously detected inversion signature against *O* females, and transcripts expressed in at least one adult individual of both *I. elegans* and *I. senegalensis*. Transcripts plotted in black are present in both the *A* and *O* assemblies, while transcripts in blue are located in genomic regions that are unique to the *A* haplotype or are shared between *A* and *I* but not the *O* allele. **b** Functional annotations and sex– and morph-specific expression of transcripts. Square fill indicates whether transcript expression was detected in each group. RNAseq data for *I. elegans* comes from whole-thorax samples from sexually immature and sexually mature wild-caught adults (n = 3 females of each morph and 3 males). RNAseq data for *I. senegalensis* comes from a recent study in which the abdomen, head, thorax, and wings were sampled in two females of each morph and two males (one individual of each group sampled upon emergence and one sampled after two days).

Three transcripts (from two predicted genes) in the morph locus are expressed in *A* females of *I. senegalensis*, and in *A* and *I* females of *I. elegans*, but never in *O* or *O*-like females (Fig. 6b). Only one of these gene models (Afem.4094) could be functionally annotated, and appears to encode a Long Interspaced Nuclear Element (LINE) retrotransposon in the clade Jockey (Supporting Text 6). This gene also exhibited expression changes in *I* females that reflect their colour development trajectory of initial resemblance to *A* females, followed by an overall appearance similar to *O* females upon sexual maturation (Supporting Text 6). Notably, *RepeatModeler* and *RepeatMasker* detected signatures of the Jockey family at the same locus as the mapping locations of the *A* reads that had suggested a propagation of TEs in our SV analyses (Fig. 6a; Extended Data Fig. 3). Thus, these results further support that TEs are responsible for the evolution and expansion of the male-mimicry allele.

We also identified three gene models that are shared by all haplotypes and expressed in both species. The three predicted genes encode zinc-finger domain proteins (Fig. 6b; Supporting Text 6), which are known to participate in transcriptional regulation^39^. However, we found no conclusive evidence of differential expression, nor evidence of non-synonymous substitutions between morphs shared by both *I. elegans* and *I. senegalensis* (Supporting Text 6). While we see genes of a potentially regulatory function reside in the morph locus, understanding their role in morph development will likely require higher temporal and spatial resolution of gene expression data.

## Discussion

Sexual dimorphism, where males and females have markedly distinct colour patterns, has led to multiple evolutionary origins of female-limited polymorphisms and potential male-mimicry in *Ischnura* damselflies^28^. Here, we present a first genomic glance into how these morphs evolve, setting the stage for future functional work to unravel the reversal of sexual phenotypes in damselfly sexual mimicry. Male mimicry in the Common Bluetail is controlled by a single genomic region in chromosome 13 (Fig. 2; 3). Our data suggests that this morph locus likely evolved with the accumulation of novel and potentially TE-derived sequences in the male mimicry haplotype (Fig. 4), which is shared by male-mimicking females of species diverging more than 5 Ma (Fig. 5). More recently, a rare recombination event involving part of the novel *A* genomic content has triggered the origin of a third female morph (Fig. 4), which shares its sexually immature colouration and patterning with *A* females, and shares its sexually mature overall appearance with *O* females^27^. The morph locus contains a handful of genes, some of which may have evolved with TE propagation in the *A* haplotype, and are therefore absent from *O* individuals (Fig. 6). However, existing annotations provide only a hint on how these genes may influence morph development. Our results thus echo recent calls for a broader application of functional validation tools, in order to understand how lineage-specific genes contribute to phenotypic variation in natural populations^40^.

This study underscores two increasingly recognized insights in evolutionary genomics. First, there is mounting evidence that structural variants abound in natural populations and often underpin complex and ecologically relevant phenotypic variation^41^, such as discrete phenotypic polymorphisms^10,13,15,20^. Nonetheless, traditional GWAS approaches based on SNPs can easily miss structural variants, as these approaches are contingent on the genomic content of the reference assembly^42^. Among other novel approaches to tackle this problem^42^, a reference-free *k*-mer based GWAS, as implemented here, is a powerful method to identify variation in genomic content and sequence, especially when the genomic architecture of the trait of interest is initially unknown^35^. In this study, we did not know *a priori* which of the three morphs, if any, would harbour unique genomic content. Had we ignored differences in genomic content between morphs and based our GWAS analysis solely upon the DToL (*O*) reference assembly, we would have failed to identify SNPs between *I* and *O* morphs (Extended Data Fig. 8), and the origin of *I* females via a translocation of *A* content would have been obscured.

Second, a role for TEs in creating novel and even adaptive phenotypic variation is increasingly being recognized^43,44^. Here, we found that a ∼400 kb region of unique genomic content, possibly driven by LINE transposition is associated with the male-mimicry phenotype in at least two species of *Ischnura* damselflies. TE activity can contribute to phenotypic evolution by multiple mechanisms. For instance, TEs may modify the regulatory environment of genes in their vicinity, by altering methylation^45^ and chromatin conformation patterns^46^, or by providing novel cis-regulatory elements^47^. The male-mimicry region in *I. elegans* is located between two coding genes with putative DNA-binding domains, and which may thus act as transcription factors. However, our expression data does not provide unequivocal support for differential regulation of either of these genes between female morphs. Importantly, currently available expression data comes from adult specimens, as female morphs are not visually discernible in aquatic nymphs. Yet, the key developmental differences that produce the adult morphs are likely directed by regulatory variation during earlier developmental stages. Now that the morph locus has been identified, future work can address differential gene expression at more relevant developmental stages, before colour differences between morphs become apparent.

TEs can also contribute to phenotypic evolution if they become domesticated, for example, when TE-encoded proteins are remodeled through evolutionary change to perform adaptive host functions^48^. We found two transcripts located in *A* specific or *A/I* specific regions that are likely derived from LINE retrotransposons and are actively expressed in the genomes that harbour them (Fig. 6b). It is therefore possible that these transcripts participate in the development of adult colour patterns, which are initially more similar between *A* and *I* females than between either of these morphs and *O* females^27,29^. Yet, functional work on these transcripts is required to ascertain their role in morph determination. Finally, TEs can become sources of novel small regulatory RNAs which play important regulatory roles^49^, including in insect sex determination^50^. Thus, future work should also address non-coding RNA expression and function in the morph locus.

Our results also provide molecular evidence for previous insights, gained by alternative research approaches, on the micro– and macroevolution of female-limited colour polymorphisms. A wealth of population data in Southern Sweden has shown that female-morph frequencies are maintained by balancing selection, as they fluctuate less than expected due to genetic drift^31^.

Behavioural and field experimental studies indicate that such balancing selection on female morphs is mediated by negative frequency-dependent male harassment^51,52^. We add to these earlier results, by showing a molecular signature consistent with balancing selection in the genomic region where *A* females differ from both of the non-mimicking morphs. Sexual conflict is expected to have profound effects on genome evolution, but there are few examples of sexually-antagonistic traits with a known genetic basis, in which predictions about these genomic effects can be tested^53,54^. Here, the signature of balancing selection on the morph locus matches the expectation of inter-sexual conflict resulting in negative-frequency dependent selection and maintaining alternative morph alleles over long periods.

Similarly, a recent comparative study based on phenotypic and phylogenetic data inferred a single evolutionary origin of the male-mimicking morph shared by *I. elegans* and *I. senegalensis*^28^. Our present results strongly support this common origin. This is because *A* females in both species share unique genomic content, including associated transcripts, and an inversion signature against the ancestral *O* morph (Fig. 5; Extended Data Fig. 7). Nonetheless, these data are consistent with both an ancestral polymorphism and ancestral introgression being responsible for the spread of male mimicry across the clade. A potential role for introgression in the evolution of male mimicry is also suggested by rampant hybridization between *I. elegans* and its closest relatives^55^, and by the fact that *I. elegans* and *I. senegalensis* can hybridize millions of years after their divergence, at least in laboratory settings^56^. The identification of the morph locus in *I. elegans*, enables future comparative genomics studies to disentangle the relative roles of long-term balancing selection and introgression in shaping the widespread phylogenetic distribution of female-limited polymorphisms in *Ischnura* damselflies.

Finally, our results open up new lines of inquiry on how the genomic architecture and chromosomal context of the female polymorphism may influence its evolutionary dynamics. Our data is consistent with the evolution of a third morph due to an ectopic recombination event that translocated genomic content from the *A* haplotype back into an *O* background. Ectopic recombination can occur when TE propagation generates homologous regions in different genomic locations^57,58^, and may be facilitated by the excess of TE content in chromosome 13 (Exteded Data Fig. 4). The male reproductive morphs in the Ruff (*Calidris pugnax*) are one of few previous examples of a novel phenotypic morph arising via recombination between two pre-existing morph haplotypes^10^. In pond damselflies, female polymorphisms with three or more female morphs are not uncommon, and in some cases female morphs exhibit graded resemblance to males^59^. It is therefore possible that recombination, even if rare, has repeatedly generated diversity in damselfly female morphs.

While recombination might have had a role in generating the the novel *I* morph, we observe reduced recombination over the morph locus in comparison to the rest of the genome of *I. elegans* (Extended Data Fig. 5). However, this reduction in recombination is not limited to the morph locus and its vicinity, but rather pervasive across chromosome 13 (Extended Data Fig. 5). This unexpected finding suggests two alternative causal scenarios. First, selection for reduced recombination at the morph locus, following the origin of sexual mimicry, could have spilled over chromosome 13, facilitating a subsequent accumulation of TEs and further reducing recombination^60^. Second, TE enrichment and reduced recombination may have preceded the evolution of female morphs, and facilitated the establishment and maintenance of the female polymorphism by balancing selection.

Both historical scenarios are compatible with a morph locus reminiscent of a supergene, which is defined by tight genetic linkage of multiple functional loci^61^. However, an alternative and parsimonious explanation is that the novel sequences in *A* and *I* females and their flanking genes may not code for anything important for the male-mimicking phenotype as such, but simply disrupt a region of chromosome 13 that is required for the development of ancestral sexual differentiation. The observation that *I* females carry part of the sequence that originated in *A* in a different location of the scaffold (Fig. 4b), and still develop some male-like characters (e.g. black thoracic stripes), could come about if insertions anywhere on a larger chromosomal region disrupt female suppression of the male phenotype, although with variable efficacy depending on the exact location or insertion size.

### Concluding Remarks

Recent years have witnessed an explosion of studies uncovering the loci behind complex phenotypic polymorphisms in various species. An emerging outlook is that not all polymorphisms are created equal, with some governed by massive chromosomal rearrangements^15–17^, and others by a handful of regulatory sites^11,12,18^. Our results contribute to this growing field by uncovering a single causal locus, that features structural variation and morph-specific transcripts, in the female-limited morphs of *Ischnura* damselflies. These morphs not only differ in numerous morphological and life-history traits^32,62,63^ and gene expression profiles^24,25^, but they include a male mimic that is maintained by balancing frequency-dependent selection. Our findings enable future studies on the developmental basis of such male mimicry, with consequences for a broader understanding of the evolutionary dynamics of sexual dimorphism and the cross-sexual transfer of trait expression^64^.

## Methods

### Ischnura elegans samples

Samples for morph-specific genome assemblies of *I. elegans* were obtained from F1 individuals with genotypes *Ao*, *Io* and *oo* (one adult female of each genotype). In June 2019, recently-mated *O* females were captured in field populations in Southern Sweden. These females oviposited in the lab within 48 h, and their eggs were then released into outdoor cattle tanks seeded with *Daphnia* and covered with synthetic mesh. Larvae thus developed under normal field conditions and emerged as adults during the Summer of 2020. Emerging females were kept in outdoor enclosures until completion of adult colour development^25,65^. Fully mature females were phenotyped, collected in liquid nitrogen and kept at –80 °C. Because all of these females carry a copy of the most recessive allele *o*, individuals of the *A* and *I* morph are heterozygous, with genotypes *Ao* and *Io*, respectively.

A total of 19 resequencing samples of each female morph of *I. elegans* were also collected from local populations in Southern Sweden, within a 40 x 40 km area (Table S1). Samples were submerged in 95% ethanol and stored in a –20 °C freezer until extraction. Additionally, 24 individuals (six adult females of each morph and six males) were collected for RNAseq analysis in a natural field population (Bunkeflostrand) in Southern Sweden, in early July 2019. These samples were transported on carbonated ice and stored in –80 °C until extraction.

### Ischnura senegalensis samples

Adults of *I. senegalensis* (30 adult females of each morph) were collected for pool sequencing from a population on Okinawa Island in Japan (26.148N, 127.795E) in May 2016. Samples were visually determined to sex and morph and stored in 99% ethanol until extraction. Samples for morph-specific genome assemblies of *I. senegalensis* were obtained from a population in Clementi Forest, Singapore (1.33N, 103.78E). Because the *A* allele is recessive in *I. senegalensis*, all females with the *A* phentoype are homozygous. To obtain a homozygous *O*-like sample, we developed primers (forward: CGCGGTATGATATGGTCCGA, reverse: GGCTGCTTACACCAATGCAA) for an *A*-specific sequence that is shared by *A* females of the two species (318,131 – 318,213 bp on the *A* haplotype of *I. elegans*). We used the mapped pool-seq data to identify fixed SNPs between species and tailor the primer sequences accordingly. We then tested the primers in 20 *A* females of *I. senegalensis* using a 328 bp fragment of the Histone H3 gene (forward: ATGGCTCGTACCAAGCAGACGGC, reverse: ATATCCTTGGGCATGATGGTGAC)^66^ as a positive control for the PCR reaction. Once validated, we utilize these primers to identify *O*-like females lacking the *A* allele and selected one of these samples for whole genome sequencing.

### DNA extraction, library preparation and sequencing

High molecular weight (HMW) DNA was extracted from one *I. elegans* female of each genotype (*Ao*, *Io*, *oo*), using the Nanobind® Tissue Big Extraction Kit (Cat. No. NB-900-701-01, Circulomics Inc. (PacBio), MD, USA). HMW DNA was isolated from homozygous females of each morph of *I. senegalensis*, using the Monarch® HMW DNA Extraction Kit for Tissue (Cat. No. T3060S, New England BioLabs Inc., MA, USA). DNA from resequencing samples was isolated using either a modified protocol for the DNeasy Blood and Tissue Kit (Cat. No. 19053, Qiagen, Germany) or the KingFisher Cell and Tissue DNA Kit (Cat no. N11997, ThermoFisher Scientific). *I. senegalensis* DNA was extracted from muscle tissues in thoraxes using Maxwell® 16 LEV Plant DNA Kit (Cat. No. AS1420, Promega, WI, USA). Details on extraction and library preparation protocols are provided in the Supporting Text 1.

Sequencing libraries were constructed from each HMW DNA sample for the Nanopore LSK-110 ligation kit (Oxford Nanopore Technologies, UK). Adapter ligation and sequencing of *I. elegans* samples were carried out at the Uppsala Genome Centre (NGI), hosted by SciLife Lab. Each sample was sequenced on a PromethION R10.4 with 1 nuclease wash and two library loadings. Library preparation and sequencing of *I. senegalensis* samples were carried out by the Integrated Genomics Platform, Genome Institute of Singapore (GIS), A-STAR, Singapore. Each sample was sequenced on a PromethION R9.4 flow cell, with 2 nuclease washes and three library loadings.

### RNA extraction and sequencing

Whole-thorax samples were grounded into a fine powder using a TissueLyser and used as input for the Spectrum^TM^ Plant Total RNA Kit (Cat. No. STRN50, Sigma Aldrich, MO, USA), including DNase I treatment (Cat. No. DNASE10, Sigma Aldrich, MO, USA). Library preparation and sequencing were performed by SciLife Lab at the Uppsala Genome Centre (NGI). Sequencing libraries were prepared from 300 ng of RNA, using the TrueSeq stranded mRNA library preparation kit (Cat. No. 20020595, Illumina Inc., CA, USA) including polyA selection and unique dual indexing (Cat. No. 20022371, Illumina Inc., CA, USA), according to the manufacturer’s protocol. Sequencing was performed on the Illumina NovaSeq 6000 SP flowcell with paired-end reads of 150 bp.

### De novo genome assembly

Bases in raw ONT reads from *I. elegans* were called using *Guppy* v 4.0.11 (*Ao* and *Io* data) and *Guppy* v 5.0.11 (*oo* data) (https://nanoporetech.com/). Low quality reads (qscore < 7 for v 4.0.11 and < 10 for v 5.0.11) were subsequently discarded. High quality reads were assembled using the *Shasta* long-read assembler v 0.7.0^67^. Each assembly was conducted under four different configuration schemes, which modified the June 2020 Nanopore configuration file (https://github.com/chanzuckerberg/shasta/blob/master/conf/Nanopore-Jun2020.conf) in alternative ways (Table S3). Assembly metrics were compared among Shasta configurations for each morph using *AsmQC*^68^ (https://sourceforge.net/projects/amos/) and the *stats.sh* script in the *BBTools* suite (https://sourceforge.net/projects/bbmap). The assembly with greater contiguity (i.e. highest contig N50, highest average contig length and highest percentage of the main genome in scaffolds > 50 kb) was selected for polishing and downstream analyses.

Bases in raw ONT reads from *I. senegalensis* samples were called using *Guppy* v 6.1.5. Reads with quality score < 7 were subsequently discarded. High quality reads were assembled using the *Shasta* long-read assembler v 0.7.0^67^ and the configuration file T2 (Table S3), which was also selected for the *Io* and *oo* assemblies of *I. elegans*.

Morph specific assemblies of *I. elegans* were first polished using the ONT reads mapped back to their respective assembly with *minimap2* v 2.22-r1110^69^, and the *PEPPER-Margin-DeepVariant* pipeline r0.4^70^. Alternative haplotypes were subsequently filtered using *purge_dups* v 0.0.3^71^, to produce a single haploid genome assembly for each sample. The *I. elegans* draft assemblies were then polished with short read data from one resequencing sample (TE-2564-SwD172_S37, Table S1), using the *POLCA* tool in *MaSuRCA* v 4.0.4^72^. For every draft and final assembly of *I. elegans*, we computed quality metrics as mentioned above and assessed the completeness of conserved insect genes using *BUSCO* v 5.0.0^73^ and the “insecta_odb10” database (Fig. S1). For *I. senegalensis*, we report quality metrics of the final assemblies (Fig. S2).

### Scaffolding with the Darwin Tree of Life super assembly

During the course of this study, a chromosome-level genome of *I. elegans* was assembled by the Darwin Tree of Life (DToL) Project^34^, based on long-read (PacBio) and short-read (Illumina) data, as well as Hi-C (Illumina) chromatin interaction data. 99.5% of the total length of this assembly is distributed across 14 chromosomes, one of which (no. 13) is fragmented and divided into a main assembly and five unlocalized scaffolds.

We used RagTag v 2.10^74^ to scaffold each our morph-specific assemblies based on the DToL reference (Supporting Text 2). Scaffolding was conducted using the *nucmer* v 4.0.0^75^ aligner and default RagTag options. Morph-specific scaffolded genomes were also aligned to each other using *nucmer* and a minimum cluster length of 100 bp. Alignments were then filtered to preserve only the longest alignments in both reference and query sequences, and alignments of at least 5 kb. These assembly alignments were then used to visualize synteny patterns across morphs, in the region uncovered in our association analyses (Extended Data Fig. 1), using the package *RIdeogram* v 0.2.2^76^ in *R* v 4.2.2^77^.

### Reference-based (SNP) GWAS

We first investigated genomic divergence between morphs using a standard GWAS approach based on SNPs (Extended Data Fig. 1). Initially, we conducted preliminary analyses using different morph assemblies as mapping reference. Once the *A*-specific genomic region was confirmed, we designated the *A* assembly as the mapping reference for the main analyses. Short-read data were mapped using *bwa-mem* v 0.7.17^78^. Optical and PCR duplicates were then flagged in the unfiltered *bam* files using *GATK* v 4.2.0.0^79^. Variant calling, filtering and sorting were conducted using bcftools v 1.12^80^, excluding the flagged reads. We retained only variant sites with mapping quality > 20, genotype quality > 30 and minor allele frequency > 0.02 (i.e. the variant is present in more than one sample). To avoid highly repetitive content, we filtered variants that had a combined depth across samples > 1360 (equivalent to all samples having ∼ 50% higher than average coverage), and variants located in sites annotated as repetitive in either *RepeatMasker* v 1.0.93^81^ or *Red* v 0.0.1^82^. The final variant calling file was analysed in pairwise comparisons (*A* vs *O*, *A* vs *I*, *I* vs *O*) using *PLINK* v 1.9^83^ (http://pngu.mgh.harvard.edu/purcell/plink/). We report the –Log_10_ of P-values for SNP associations in these pairwise comparisons.

### Reference free (k-mer) GWAS

We created a list of all *k*-mers of length 31 in the short-read data (19 females per morph, Extended Data Fig. 1) following Voichek and Weigel^35^, and counting *k*-mers in each sample using *KMC* v 3.1.0^84^. The *k*-mer list was filtered by the minor allele count; *k*-mers that appeared in less than 5 individuals were excluded. *k*-mers were also filtered by percent canonized (i.e. the percent of samples for which the reverse complement of the *k*-mer was also present). If at least 20% of the samples including a given *k*-mer contained its canonized form, the *k*-mer was kept in the list. The *k*-mer list was then used to create a table recording the presence or absence of each *k*-mer in each sample. A kinship matrix for all samples was calculated from this *k*-mer table, and was converted to a *PLINK*^83^ binary file, where the presence or absence of each *k*-mer is coded as two homozygous variants. In this step, we further filtered the *k*-mers with a minor allele frequency below 5%.

Because a single variant, be it a SNP or SV, will likely be captured by multiple *k*-mers, significance testing of *k*-mer associations requires a method to control for the non-independence of overlapping *k*-mers. We followed the approach developed by Voichek and Weigel^35^, which uses a linear mixed model (LMM) genome-wide association analysis implemented in *GEMMA* v 0.98.5^85^, and computes P-value thresholds for associated *k*-mers based on phenotype permutations. We thus report *k*-mers below the 5% false-positive threshold as *k*-mers significantly associated with the female-polymorphism in *I. elegans*. We conducted three *k*-mers based GWAS: 1) comparing male mimics to the putatively ancestral female morph (*A* vs *O*), 2) comparing male mimics to the most derive female morph (*A* vs *I*), and 3) comparing both derived female morphs (*A* and *I*) to the ancestral *O* females. For every analysis, we then mapped the significant *k*-mers to all reference genomes using *Blast* v 2.22.28^86^ for short sequences, and removed alignments that were below 100% identity and below full-length. The mapped *k*-mers thus indicate the proportion of relevant genomic content present in each morph and how this content is distributed across each genome (Extended Data Table 1).

### Read depth analysis

To validate the *k*-mer GWAS results of unique genomic content in *A*-females relative to both *I*– and *O*-females, we plotted read-depth across our region of interest (the unlocalized scaffold 2 of chromosome 13, see Results) in the *A* assembly (Extended Data Fig. 1). Short-read data (19 samples per morph) were mapped to the assembly with *bwa-mem* v 0.7.17^78^ and reads with mapping score < 20 were filtered, using *Samtools* v 1.14^87^. Long-read data (one sample per morph) were also mapped to the assembly using *minimap2* v 2.22-r1110^69^, and quality filtering was conducted as above. Read depth was then averaged for each sample across 500 bp, non–overlapping windows using *mosdepth* v 0.2.8^88^. We also annotated repetitive content in the reference genome using *RepeatMasker* v 1.0.93^81^ and *Red* v 0.0.1^82^, and filtered windows with more than 10% repetitive content under either method.

To account for differences in overall coverage between samples, we conducted the same procedure on a large (∼15 mb) non-candidate region in chromosome 11 and calculated a “background read depth” as the mean read depth across the non-repetitive windows of this region. We then expressed read-depth in the candidate region as a proportion of the background read depth. Values around 1 thus indicate that a sample is homozygous for the presence of the sequence in a window. Values around 0.5 suggest that the sample only has one copy of this sequence in its diploid genome (i.e. it is heterozygous). Finally, values of 0 imply that the 500 bp reference sequence is not present in the sample (i.e. the window is part of an insertion or deletion).

We also investigated read-depth coverage on the *I* assembly, specifically across the region that was identified in the *k*-mer based GWAS as capturing content that differentiated both *A* and *I* females from *O* females (Fig. 3b, Extended Data Fig. 1). To do so, we followed the same strategy as a above, except here we used a 15 mb region from chromosome 1 to estimate background read depth.

### Population genetics

We investigated the evolutionary consequences of morph divergence by estimating between-morph Fst and population-wide Tajima’s D and nucleotide diversity (π). For these analyses, we used the *A* assembly as mapping reference and the same variant calling approach as described for the SNP based GWAS, but applied different filtering criteria (Extended Data Fig. 1).

Specifically, invariant sites were retained and we only filtered sites with mapping quality score < 20 and combined depth across samples > 1360 (equivalent to ∼50% excess coverage in all samples). Fst and (π) were estimated in *pixy* v 1.2.5^89^ across 30 kb windows. Fst was computed using the *hudson* estimator^90^. Negative Fst values were converted to zero for plotting. Tajima’s D was estimated across 30 kb using *vcftools* v 0.1.17^91^. In all analyses, windows with > 10% repetitive content according to either *RepeatMasker* v 1.0.93^81^ or *Red* v 0.0.1^82^ annotation were excluded.

### Structural variants

We used two complimentary approaches to identify SVs overlapping the genomic region uncovered by both *k*-mer-based and SNP-based GWAS. First, we mapped the raw data from each long-read sample to the assemblies of alternative morphs (e.g. *Ao* data mapped to *Io* and *oo* assemblies), and called SVs using *Sniffles* v 1.0.10^92^ (Extended Data Fig. 1). These SV calls may represent fixed differences between morphs, within-morph polymorphisms, or products of assembly error. We therefore used *SamPlot* v 1.3.0^93^ and our short-read samples (n = 19 per morph) to validate morph-specific SV calls (Extended Data Fig. 1). *Samplot* identifies and plots reads with discordant alignments, which can result from specific types of SVs. For example, if *Sniffles* called a 10 kb deletion in the *Ao* and *Io* long-read samples relative to the *oo* assembly, we then constructed a Samplot for this region using short-read data, and expected to find support for such deletion in *I* and *A* samples, but not in *O* samples. We complemented this validation approach with a scan of the region of interest in each assembly, in windows of 250 and 500 kb, again using *Samplot* and the short-read data. If a SV appeared to be supported by the majority of short-read samples from an alternative morph, we zoomed in this SV and recorded the number of samples supporting the call in each morph.

### Linkage disequilibrium and transposable elements

To estimate linkage disequilibrium (LD), we used the same variant calling file as for the SNP based GWAS, which included only variant sites and was filtered by mapping quality, genotyping quality, minimum allele frequency, and read depth, as described above (Extended Data Fig. 1).

The file was downsampled to one variant every 100th using vcftools v 0.1.17^91^, prior to LD estimation. We estimated LD using *PLINK* v 1.9^83^, and recorded R^2^ values > 0.05 for pairs up to 15 mb apart or with 10,000 or fewer variants between them. We estimated LD for the unlocalized scaffold 2 of chromosome 13, which contains the morph loci and is about ∼ 15 mb in the *A* assembly. For comparison, we also estimated LD across the first 15 mb of the fully assembled chromosomes (1-12 and X), the main scaffold of chromosome 13, and the unlocalized scaffolds 1, 3 and 4 of chromosome 13.

We used the TE annotations from *RepeatModeler* v 2.0.1 *RepeatMasker* v 1.0.93^81^ and “One code to find them all”^94^ to quantify TE coverage in chromosome 13 in comparison to the rest of the genome. We divided each chromosome into 1.5 mb windows, and computed the proportion of each window covered by each TE family.

### Evidence of a trans-species polymorphism

We used pool-seq data from the closely related Tropical Bluetail damselfly (*Ischnura senegalensis*) to determine whether male-mimicry has a shared genetic basis in the two species (Extended Data Fig. 1). First, we aligned the short-read data from the the two *I. senegalensis* pools (*A* and *O*-like) to the *A* morph assembly of *I. elegans* using *bwa-mem* v 0.7.17^78^, and filtered reads with mapping score < 20, using *Samtools* v 1.14^87^. We then quantified read depth as for the *I. elegans* resequencing data (see *Read depth analysis* above). To confirm that the higher read-depth coverage of the *A* pool is specific to the putative morph locus, we also plotted the distribution of read-depth differences between *O*-like and *A* pools across the rest of the genome and compared it to the morph locus (Supporting Text 5). Next, we determined if the ∼20 kb SV that characterizes *A* and *I* females of *I. elegans* is also present in *A* females of *I. senegalensis*. To do this, we mapped the pool-seq data to the *O* assembly of *I. elegans* as above, and scanned the region at the start of the scaffold 2 of chromosome 13 for SVs using Samplot v1.3.0^93^. Finally, we aligned the morph-specific assemblies of *I. senegalensis* to the *A* assembly of *I. elegans*, using *nucmer* v 4.0.0^75^ and preserving alignments > 500 bp and with identity > 70% (Extended Data Fig. 1). We visualized synteny patterns across the morph locus using the package *RIdeogram* v 0.2.2^76^ in *R* v 4.2.2^77^.

### Gene content and expression in the morph locus

We assembled transcripts in the *A* morph genome (Extended Data Fig. 1) to identify potential gene models unique to the *A* or *A* and *I* morphs and which would therefore be absent from the *I. elegans* reference annotation (based on the *O* haplotype). First, all raw RNAseq data from *I. elegans* samples were mapped to the *A* assembly using HISAT2 v 2.2.1^95^ and reads with mapping quality < 60 were filtered using *Samtools* v 1.19^87^. Transcripts were then assembled in *StringTie* v 2.1.4^96^ under default options, and merged into a single gtf file. Transcript abundances were quantified using this global set of transcripts as targets, and a transcript count matrix was produced using the *prepDE.py3* script provided with *StringTie*. Mapped RNAseq data from *I. senegalensis* was also used to assemble transcripts (Extended Data Fig. 1), but this time the HISAT2 assembly was guided by the annotation based on *I. elegans* data, while allowing the identification of novel transcripts. Transcript abundances were quantified as for *I. elegans*.

We analyzed differential gene expression using the package *edgeR* v 3.36^97^ in *R* v 4.2.2^77^. Transcripts with fewer than one count per million in more than three samples were filtered. Library sizes were normalized across samples using the trimmed mean of M-values method^98^, and empirical Bayes tagwise dispersion^99^ was estimated prior to pairwise expression analyses. Differential expression of genes in the morph loci was tested using two-tailed exact tests^100^, assuming negative-binomially distributed transcript counts and applying the Benjamini and Hochberg’s algorithm to control the false discovery rate (FDR)^101^.

Nucleotide sequences of all transcripts mapped to the 1.5 mb morph locus in the *A* assembly were selected. Coding sequences (CDS) in these transcripts were predicted using *Transdecoder* v 5.5.0 (https://github.com/TransDecoder/TransDecoder). Predicted CDS and peptide sequences were read from the assemblies using the genome-based coding region annotation produced with *Transdecoder* and *gffread* v 0.12.7^102^. We investigated whether any inferred peptides or transcripts were unique to *A* or *A* and *I* by comparing these sequences to the DToL reference transcriptome and proteome (based on the *O* haplotype). We then searched for homologous and annotated proteins in other taxa in the Swissprot database using *Blast* v 2.9.0^86^. We found three gene models which are protein-coding and present in both *A* and *O* females (see Results, Fig. 6).

We scanned these protein sequences for functional domains using *InterProScan*^103^ and searched for orthologous groups and functional annotations in *EggNOG* v 5.0^104^.

### Data availability

Sequencing data from this study have been submitted to the NCBI Sequence Read Archive (SRA) (https://www.ncbi.nlm.nih.gov/sra/) under BioProject PRJNA940276. For details, please see Supplementary Tables 1 and 2. Morph-specific genome assemblies and intermediate output files required to reproduce the figures in the main text and Supporting Material are available on Zenodo^105^.

### Code availability

All code necessary to reproduce the results of this study can be found on Zenodo https://doi.org/10.5281/zenodo.8304055 and Github https://github.com/bwillink/Morph-locus.

## Acknowledgements

We thank H. Dort, R.A. Steward, J. Haushofer, P. de Sessions, and the Monteiro Lab for suggestions and helpful discussions. We are also grateful to M.P. Celorio-Mancera and H.M. Low for invaluable technical support. Prof. A. Monteiro hosted B.W. at the National University of Singapore during part of the duration of this study. Computation and data handling were enabled by resources provided by the Swedish National Infrastructure for Computing (SNIC) through the Uppsala Multidisciplinary Center for Advanced Computational Science (UPPMAX), under projects 2022/6-230 and 2022/5-419 awarded to EIS. Specimens of *I. senegalensis* were collected in Singapore, under a Research Permit (NP/RP22-015b) from the National Parks Board, Singapore. This work was supported by an International Postdoc Grant from the Swedish Research Council (VR; grant no. 2019-06444 to BW). Funding was also provided by the Swedish Research Council (VR; grant no. 2017-04386 to CWW, and grant no. 2016-03356 to EIS) and by the Stina Werners Foundation (grant no. 2018-017 to EIS) and Erik Philip Sörensens Stiftelse (grant no. 2019-033 to EIS). SN was funded by a scholarship grant for Master’s students from Sven and Lily Lawski’s Foundation.

## Author contributions

**BW** conceived the study with input from **CWW**. **EIS** organized field work in the long-term population study of *I. elegans* during 2019 and 2020, planned and prepared the outdoor rearing experiments. **EIS** and **SN** collected DNA and RNA samples of *I. elegans*. **MT** and **YT** collected samples for pool sequencing of *I. senegalensis*, and **BW** collected samples of *I. senegalensis* in Singapore. **SN**, **BW**, and **KT** conducted laboratory work on *I. elegans*. **MT**, **YT**, and **BW** conducted laboratory work on *I. senegalensis*. **BW** analyzed the data with input from **CWW**, **KT**, **RC**, and **TL**. **BW** wrote the manuscript with contributions from all authors.

## Extended data Tables

**Extended Data Table 1.** Significant *k*-mers associated with morph comparisons in *I. elegans*. For each comparison (*A* vs *O*, *A* vs *I* and *A* and *I* vs *O*), we show the total number of significant *k*-mers, and the total number of significant *k*-mers that map without any mismatching position to morph-specific reference assemblies. Of the mapping *k*-mers, we then show the number located in the unlocalized scaffold 2 of chromosome 13, which includes the putative morph locus. For the DToL assembly, we show the number of significant *k*-mers mapping to both the primary assembly (capturing the *O* allele) and the haplotigs, where the haplotig RAPID_106 comprises the *A* allele (see Supporting Text 2).

| *k*-mers | *A* vs *O* | *A* vs *I* | *I* vs *O* |
| --- | --- | --- | --- |
| Total number | 568,039 | 508,031 | 85,134 |
| *A* assembly | 435,509 | 383,679 | 45,580 |
| *A* Chr 13_2 | 427,606 | 376,075 | 44,990 |
| *I* assembly | 46,733 | 1,111 | 49,093 |
| *I* Chr 13_2 | 45,819 | 375 | 48,484 |
| *O* assembly | 1,478 | 762 | 1,452 |
| *O* Chr 13_2 | 276 | 92 | 72 |
| DToL primary assembly | 915 | 756 | 676 |
| DToL Chr 13_2 | 134 | 115 | 14 |
| DToL haplotigs | 542,571 | 489,912 | 66,861 |
| DToL RAPID_106 | 539,588 | 486,943 | 65,999 |

## Extended data figure legends

**Extended Data Figure 1.**
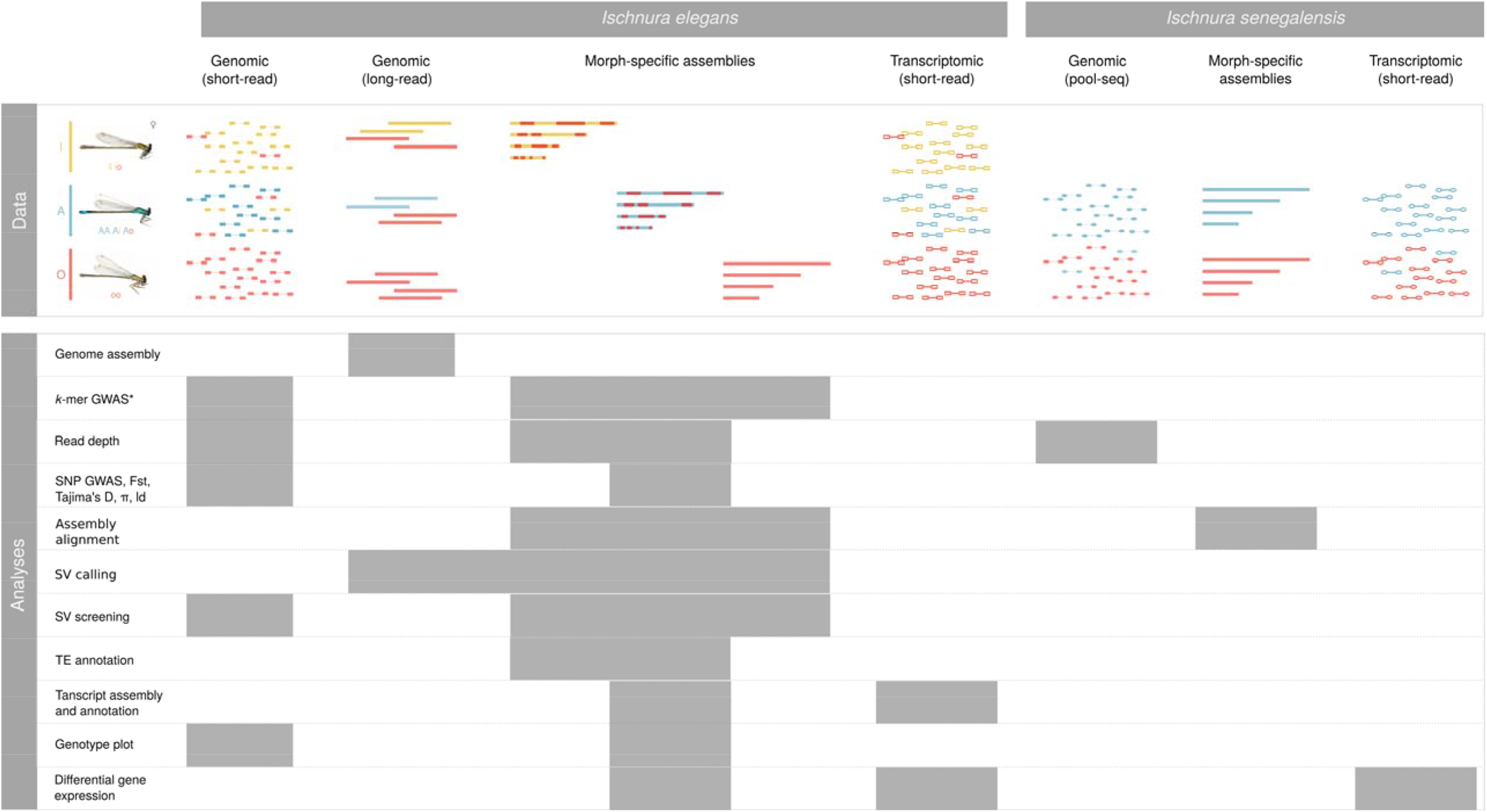
Outline of data and analyses used in this study. For our main study species *Ischnura elegans*, we obtained short-read genomic data from 19 field-caught females per morph, and long-read genomic data from three females with genotypes *Ao*, *Io*, and *oo*. The long-read samples were used to assemble morph-specific genomes, scaffolded against the Darwin Tree of Life reference assembly. We obtained whole-thorax RNAseq data from three females of each morph in both sexually immature and sexually mature colour phases (n = 2). Immature and mature males (n = 3 of each) were also sampled for whole-thorax RNAseq data. We used short-read pool-seq data (n = 30 individuals of each morph per pool) of the close relative *Ischnura senegalensis* to investigate whether the female polymorphisms in both species share a genomic basis. We also analysed expression levels of candidate genes in this species, using samples from a previously published study^38^, which produced transcriptomic data from four body parts (head, thorax, wing and abdomen) of each *A* females, *O* females and males (n = 2), sampled at adult emergence and two days thereafter. The *k*-mer based GWAS is reference-free, but significant *k*-mers were mapped to the morph-specific assemblies to determine their chromosomal context.

**Extended Data Figure 2.**
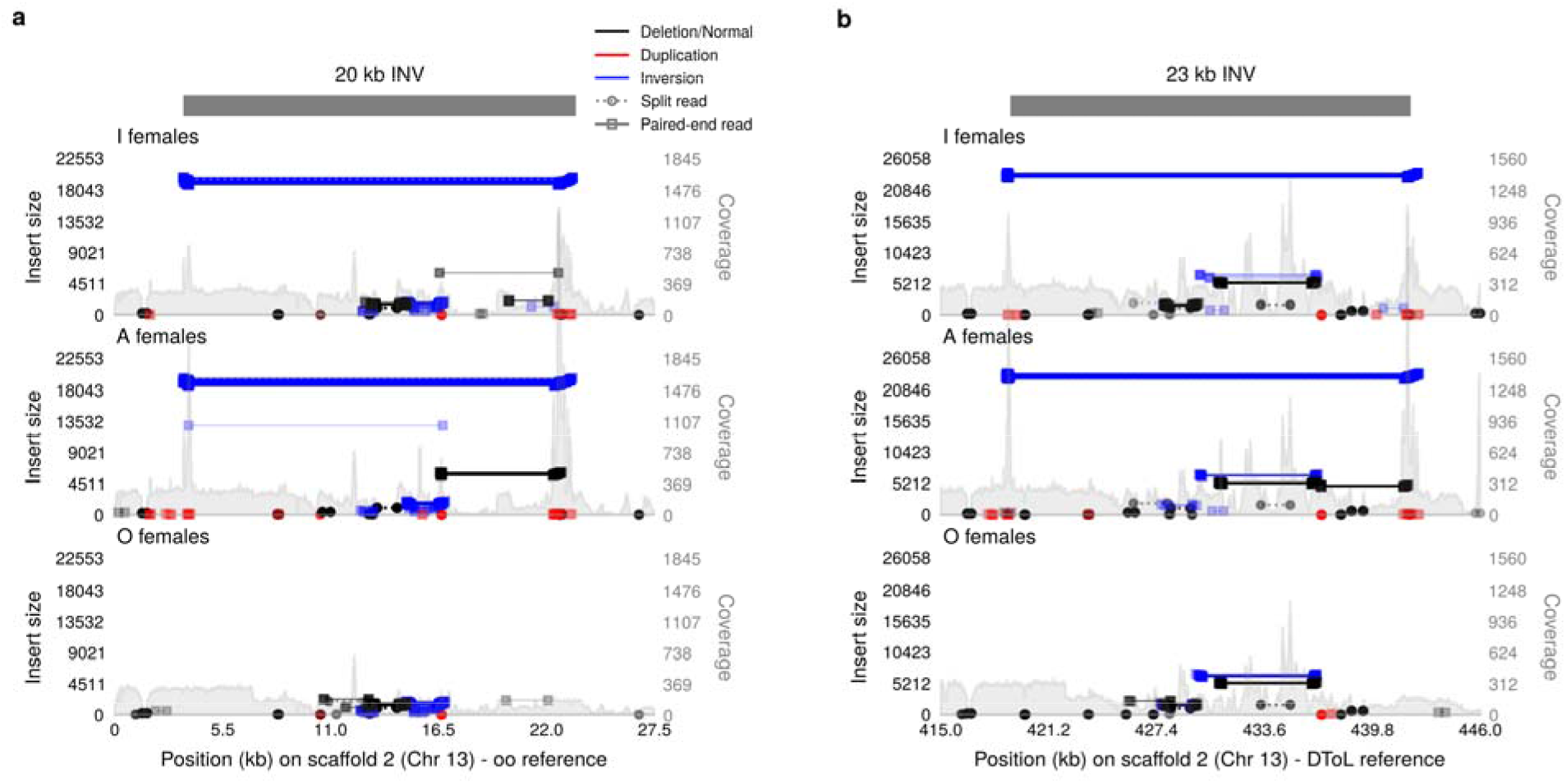
An inversion signature differentiates *A* and *I* individuals from the *O* morph. Read mapping and sample coverage at the start of the scaffold 2 of chromosome 13 in **a** our *O* assembly and **b** the DToL reference assembly, showing a signature of a ∼ 20 kb inversion in *A* and *I* samples. A single *O* sample also exhibited this signature but was excluded here for clarity (see Supporting Text 3). Note that the first 415 kb of the reference DToL assembly are absent in our scaffolded *O* assembly, and therefore the x-axis is shifted by 415 kb in **b**.

**Extended Data Figure 3.**
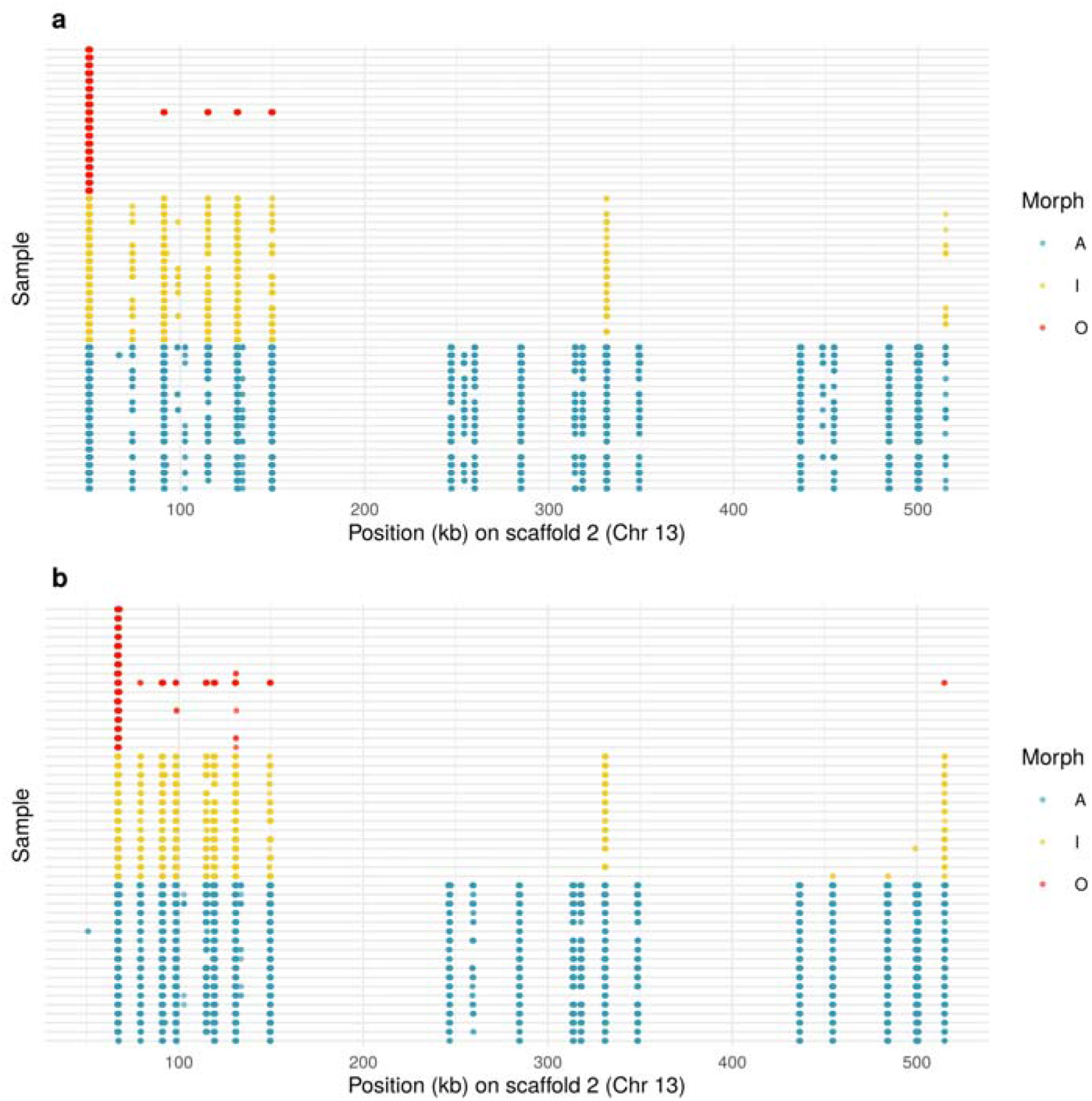
The *A* and *I* reads mapped to inversion break points on the *O* assembly (see Extended Data Fig. 2) map to multiple locations on the *A* assembly. **a** Reads from the first inversion breakpoint. **b** Reads from the second inversion breakpoint. Each row represents a sample and each circle an individual read. The x-axis corresponds to coordinates on the *A* assembly.

**Extended Data Figure 4.**
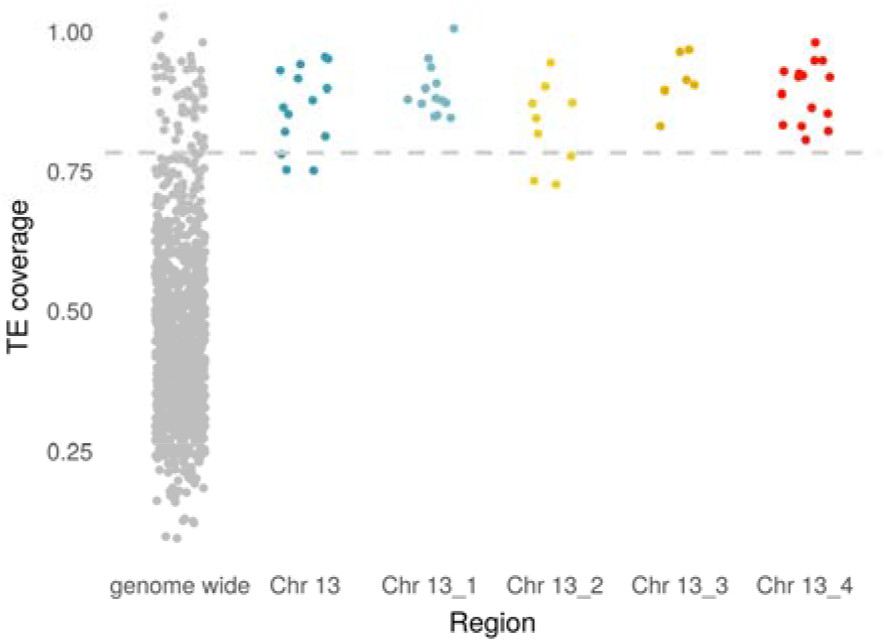
Proportion of TE content in non-overlapping 1.5 mb regions. The gray dots correspond to genomic windows outside chromosome 13. The main assembly and the unlocalized scaffolds of chromosome 13 are depicted with different colours. The dashed line marks the 95 percentile of TE coverage across all windows.

**Extended Data Figure 5.**
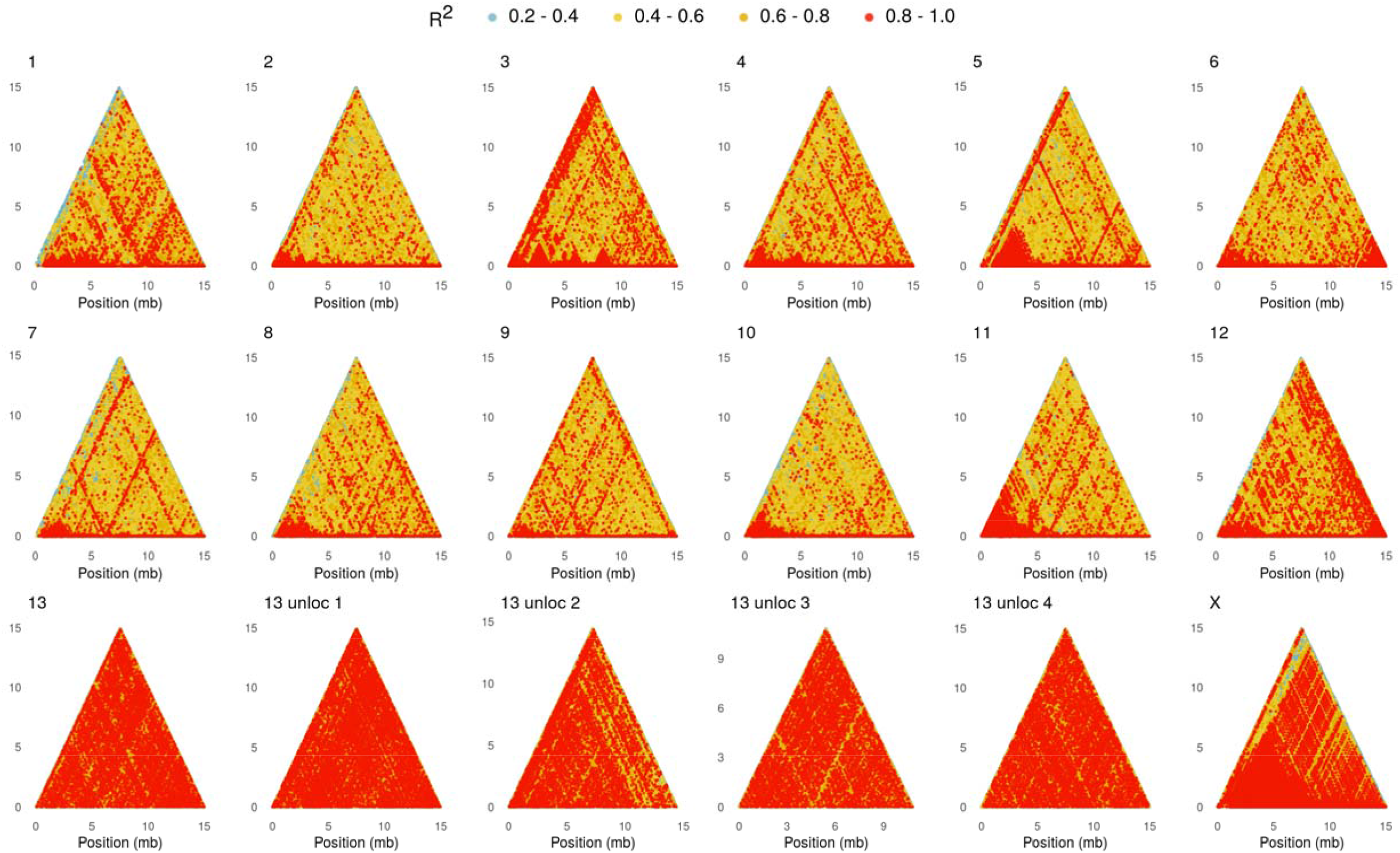
Linkage disequilibrium (LD) in the genome of *Ischnura elegans*. LD estimates are shown for the first 15 mb of each chromosome and all unlocalized scaffolds of chromosome 13. The morph locus is found in the first ∼ 1.5 mb of the unlocalized scaffold 2 of chromosome 13, which has a total size of ∼ 15 mb. Each dot represent the square correlation coefficient (R^2^) between two variant sites on the x axis, separated by the number of sites indicated in the y axis.

**Extended Data Figure 6.**
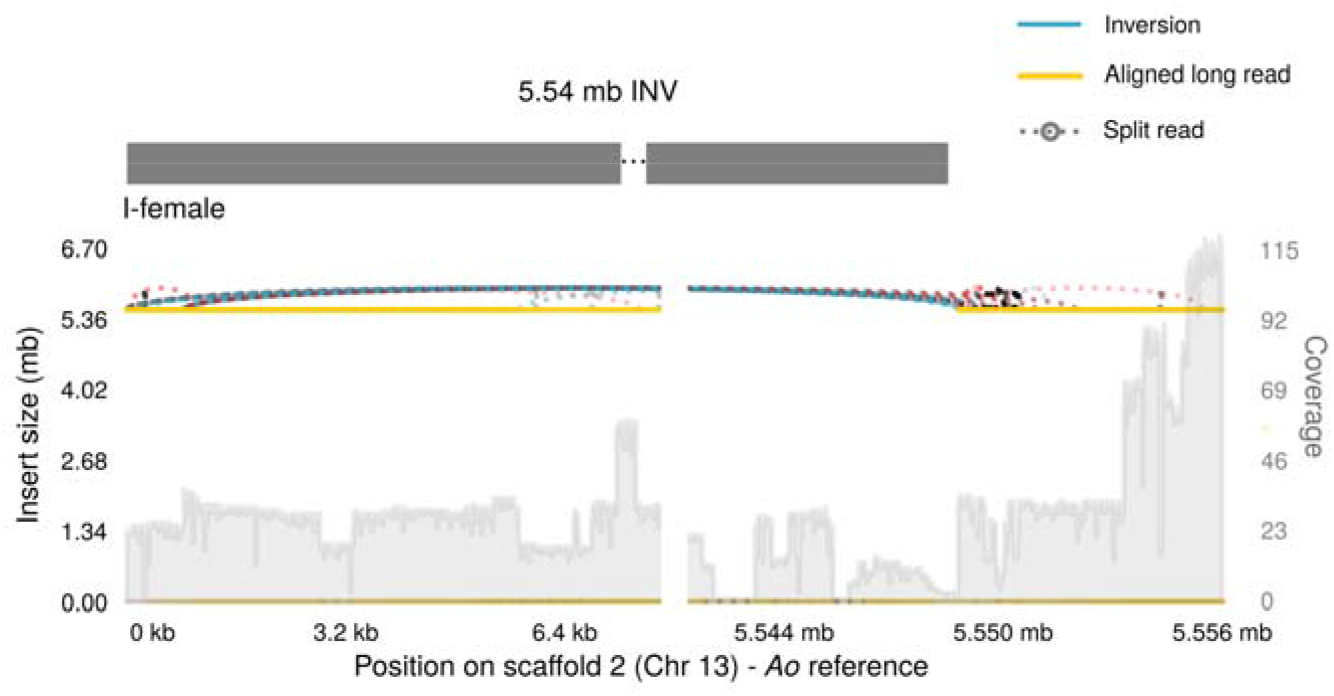
Evidence of a translocation between *A* and *I* haplotypes. Mapping and coverage of long reads from an *Io* sample across the first 5.6 mb of the unlocalized scaffold 2 of chromosome 13 in the *A* assembly, showing a signature consistent with either a 5.54 mb inversion or a translocation of inverted *A* content. Absence of morph divergence beyond ∼1.5 mb on the *A* assembly supports the translocation scenario.

**Extended Data Figure 7.**
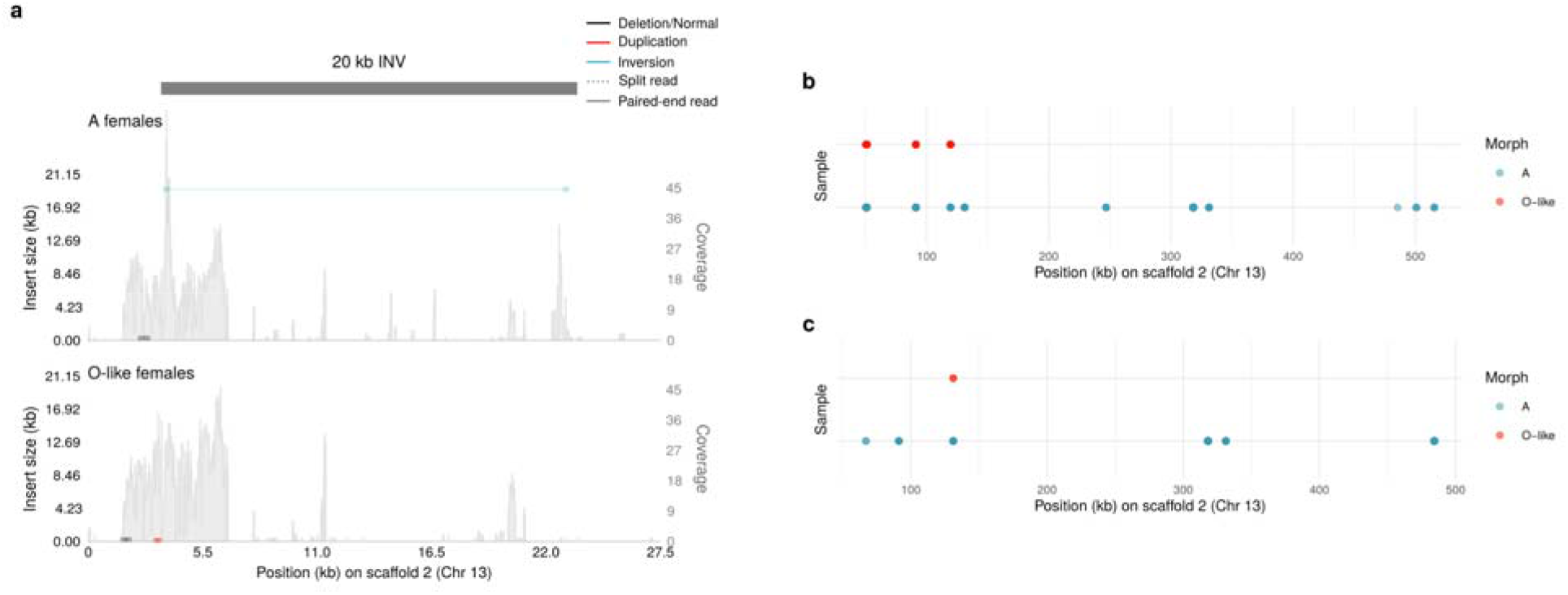
Structural variants between *A* and *O*-like females of *I. senegalensis* along the morph locus identified in *I. elegans.* **a** Read mapping and sample coverage of *I. senengalensis* pool-seq data at the start of the unlocalized scaffold 2 of chromosome 13 in the *O* assembly of *I. elegans*. The same ∼ 20 kb inversion signature is found in *A* and *I* samples of *I. elegans* (see Extended Data Fig. 2). **b-c** The *A*-pool reads mapped to break points on the *O* assembly map to multiple locations on the *A* assembly. **b** Reads from the first breakpoint. **c** Reads from the second breakpoint. Each row represents a pool of *I. senegalensis* and each circle an individual read. The x-axis corresponds to the *A* assembly of *I. elegans*.

**Extended Data Figure 8.**
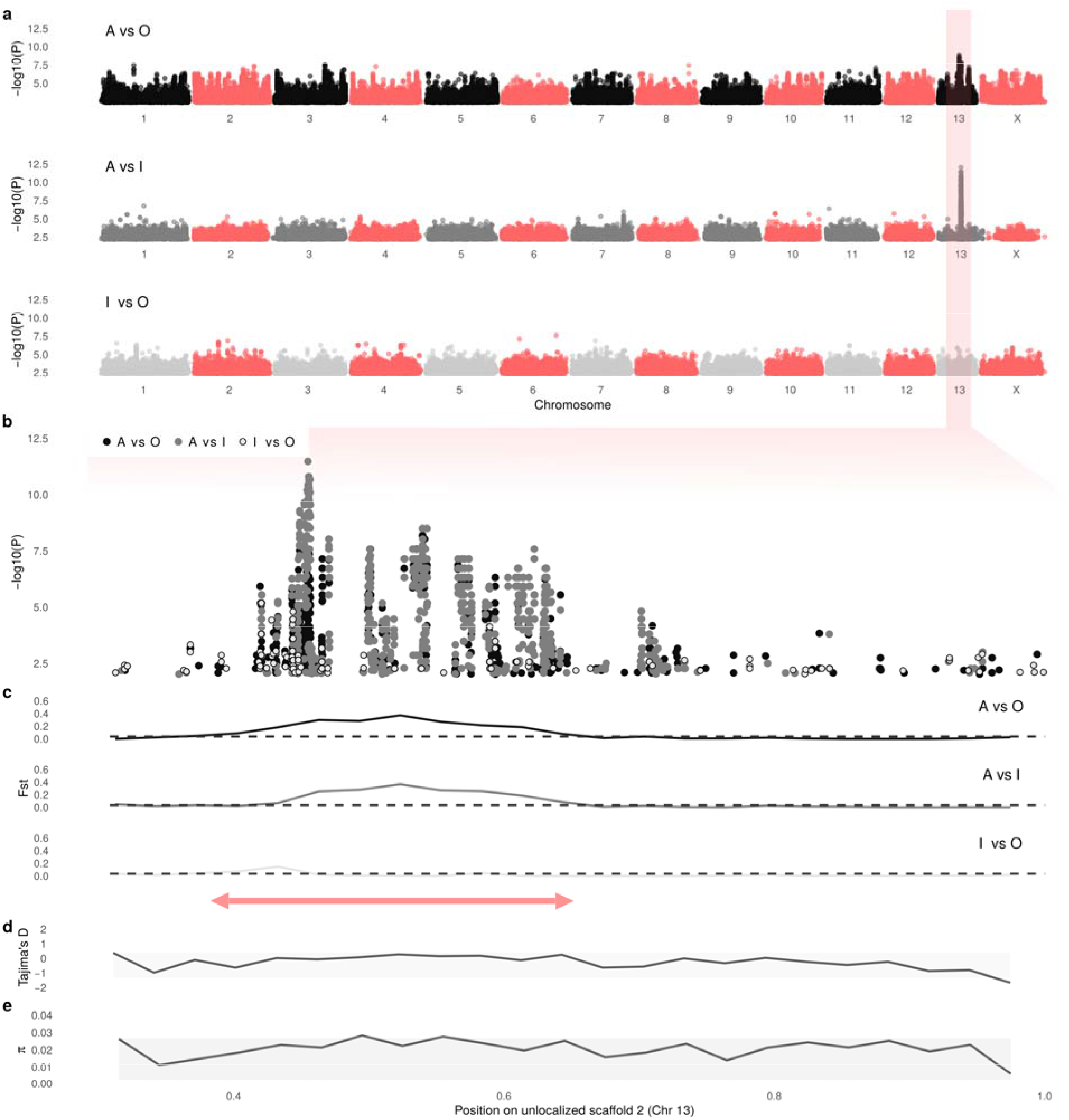
Morph divergence using the DToL assembly (*O* haplotype) as mapping reference. **a** SNP-based genome-wide associations in all pairwise analyses between morphs. Genomic DNA from 19 wild-caught females of each colour morph and of unknown genotype was extracted and sequenced for these analyses. Illumina short reads were aligned against the DToL reference assembly. **b** A closer look of the SNP associations on the unlocalized scaffold 2 of chromosome 13, which contained all highly significant SNPs. The *y* axis in **a** and **b** indicates unadjusted –Log_10_ P-values calculated from chi-squared tests. **c** Fst values averaged across 30 kb windows for the same pairwise comparisons as in the SNP based GWAS. The dashed line marks the 95 percentile of all non-zero Fst values across the entire genome. The red double arrow shows the region of elevated divergence between *A* and both *O* and *I* samples. Population-level estimates of **d** Tajima’s D, and **e** nucleotide diversity (π) averaged across 30 kb windows. The shaded area contains the 5-95 percentile of all genome-wide estimates.

