## Supplementary Texts, Figures and Tables for "The genomics and evolution of inter-sexual mimicry and female-limited polymorphisms in damselflies"

2023-09-19

[1] Department of Zoology, Stockholm University, Stockholm 106-91, Sweden

[2] Department of Biological Sciences, National University of Singapore, Singapore 117558, Singapore

[3] Department of Biology, Evolutionary Ecology Unit, Lund University, Lund 223-62, Sweden

[4] Institut Pasteur, Université Paris Cité, Sequence Bioinformatics, F-75015 Paris, France

[5] Univ. Rennes, Inria, CNRS, IRISA, Rennes F-35000, France

[6] Graduate School of Life Sciences, Tohoku University, 6-3 Aramaki, Aoba, Sendai, 980-8578, Japan

[7] Graduate School of Science, Chiba University, 1-33 Yayoi, Inage, Chiba, 263-8522, Japan

[†] Current address: Graduate School of Agriculture, Kyoto University, Oiwakecho, Kitashirakawa, Sakyoku, Kyoto, 606-8502, Japan

**References** **37**

### Sample information

**Table S1.** Resequencing samples of *Ischnura elegans*. All samples are adult female specimens caught in the field.

| Sample_ID | accession | Morph | Population |
|---|---|---|---|
| TE-2564-SwD108_S67 | SAMN33707085 | I | Bunkeflostrand |
| TE-2564-SwD136_S46 | SAMN33548694 | A | Lomma |
| TE-2564-SwD138_S47 | SAMN33548695 | A | Lomma |
| TE-2564-SwD143_S68 | SAMN33548696 | I | Bunkeflostrand |
| TE-2564-SwD149_S35 | SAMN33548697 | A | Bunkeflostrand |
| TE-2564-SwD152_S64 | SAMN33548698 | I | Borgeby |
| TE-2564-SwD153_S65 | SAMN33548699 | I | Borgeby |
| TE-2564-SwD170_S36 | SAMN33548700 | A | Bunkeflostrand |
| TE-2564-SwD172_S37 | SAMN33548701 | I | Bunkeflostrand |
| TE-2564-SwD174_S55 | SAMN33548702 | I | Genarp |
| TE-2564-SwD178_S56 | SAMN33548703 | I | Genarp |
| TE-2564-SwD19_S52 | SAMN33548704 | I | Vombs Vattenverk |
| TE-2564-SwD191_S39 | SAMN33548705 | A | Vombs Vattenverk |
| TE-2564-SwD21_S53 | SAMN33548706 | I | Vombs Vattenverk |
| TE-2564-SwD215_S62 | SAMN33548707 | I | Lomma |
| TE-2564-SwD216_S63 | SAMN33548708 | I | Lomma |
| TE-2564-SwD238_S49 | SAMN33548709 | O | Borgeby |
| TE-2564-SwD240_S66 | SAMN33548710 | I | Borgeby |
| TE-2564-SwD242_S50 | SAMN33548711 | A | Borgeby |
| TE-2564-SwD243_S51 | SAMN33548712 | A | Borgeby |
| TE-2564-SwD257_S69 | SAMN33548713 | I | Bunkeflostrand |
| TE-2564-SwD263_S57 | SAMN33548714 | I | Genarp |
| TE-2564-SwD27_S38 | SAMN33548715 | A | Vombs Vattenverk |
| TE-2564-SwD283_S40 | SAMN33548716 | A | Vombs Vattenverk |
| TE-2564-SwD288_S54 | SAMN33548717 | I | Vombs Vattenverk |
| TE-2564-SwD29_S41 | SAMN33548718 | A | Genarp |
| TE-2564-SwD308_S44 | SAMN33548719 | A | Ilstorp |

*(continued)*

| Sample_ID | accession | Morph | Population |
|---|---|---|---|
| TE-2564-SwD309_S45 | SAMN33548720 | A | Ilstorp |
| TE-2564-SwD31_S42 | SAMN33548721 | A | Genarp |
| TE-2564-SwD32_S43 | SAMN33548722 | A | Genarp |
| TE-2564-SwD51_S58 | SAMN33548723 | I | Ilstorp |
| TE-2564-SwD55_S59 | SAMN33548724 | I | Ilstorp |
| TE-2564-SwD57_S60 | SAMN33548725 | I | Ilstorp |
| TE-2564-SwD65_S48 | SAMN33548726 | A | Lomma |
| TE-2564-SwD67_S61 | SAMN33548727 | I | Lomma |
| UE-2969-57120_S90 | SAMN33548740 | O | Genarp |
| UE-2969-57123_S82 | SAMN33548732 | O | Genarp |
| UE-2969-57210_S81 | SAMN33548731 | O | Habo Gard |
| UE-2969-57263_S78 | SAMN33548728 | O | Bunkeflostrand |
| UE-2969-57302_S83 | SAMN33548733 | O | Lomma |
| UE-2969-57359_S84 | SAMN33548734 | O | Borgeby |
| UE-2969-57373_S85 | SAMN33548735 | O | Flackarp |
| UE-2969-57560_S86 | SAMN33548736 | O | Hoje A 7 |
| UE-2969-57561_S87 | SAMN33548737 | O | Hoje A 7 |
| UE-2969-57584_S97 | SAMN33548747 | A | Hoje A 14 |
| UE-2969-57846_S98 | SAMN33548748 | A | Flyinge 30 A1 |
| UE-2969-57945_S88 | SAMN33548738 | O | Vombs Vattenverk |
| UE-2969-58013_S91 | SAMN33548741 | O | IKEA |
| UE-2969-58733_S92 | SAMN33548742 | O | Lunnarp |
| UE-2969-59008_S89 | SAMN33548739 | O | Ladugardsmarken |
| UE-2969-60117_S80 | SAMN33548730 | O | Ilstorp |
| UE-2969-60681_S93 | SAMN33548743 | O | Hoje A 14 |
| UE-2969-61106_S79 | SAMN33548729 | O | Gunnesbo |
| UE-2969-61346_S99 | SAMN33548749 | A | Lunnarp |
| UE-2969-61361_S94 | SAMN33548744 | O | Lunnarp |
| UE-2969-Sw1118_S96 | SAMN33548746 | O | unknown |
| UE-2969-Sw469_S95 | SAMN33548745 | O | unknown |

**Table S2.** RNAseq samples of *Ischnura elegans*. All samples were extracted from whole-thorax tissues of adults caught in the field.

| Sample_ID | accession | Sex | Age | Morph |
|---|---|---|---|---|
| SJ-2341-SwR11_S1 | SAMN33548750 | Female | Immature | A |
| SJ-2341-SwR20_S3 | SAMN33548756 | Female | Immature | A |
| SJ-2341-SwR39_S4 | SAMN33548762 | Female | Immature | A |
| SJ-2341-SwR19_S2 | SAMN33548755 | Female | Mature | A |
| SJ-2341-SwR49_S5 | SAMN33548764 | Female | Mature | A |
| SJ-2341-SwR66_S6 | SAMN33548767 | Female | Mature | A |
| SJ-2341-SwR21_S7 | SAMN33548757 | Female | Immature | I |
| SJ-2341-SwR30_S9 | SAMN33548760 | Female | Immature | I |
| SJ-2341-SwR31_S10 | SAMN33548761 | Female | Immature | I |
| SJ-2341-SwR22_S8 | SAMN33548758 | Female | Mature | I |
| SJ-2341-SwR53_S11 | SAMN33548765 | Female | Mature | I |
| SJ-2341-SwR68_S12 | SAMN33548769 | Female | Mature | I |
| SJ-2341-SwR3_S13 | SAMN33548759 | Female | Immature | O |
| SJ-2341-SwR54_S14 | SAMN33548766 | Female | Immature | O |
| SJ-2341-SwR83_S15 | SAMN33548771 | Female | Immature | O |
| SJ-2341-SwR158_S18 | SAMN33548754 | Female | Mature | O |
| SJ-2341-SwR84_S16 | SAMN33548772 | Female | Mature | O |
| SJ-2341-SwR86_S17 | SAMN33548773 | Female | Mature | O |
| SJ-2341-SwR111_S22 | SAMN33548751 | Male | Immature | |
| SJ-2341-SwR113_S23 | SAMN33548752 | Male | Immature | |
| SJ-2341-SwR132_S24 | SAMN33548753 | Male | Immature | |
| SJ-2341-SwR4_S19 | SAMN33548763 | Male | Mature | |
| SJ-2341-SwR67_S20 | SAMN33548768 | Male | Mature | |
| SJ-2341-SwR69_S21 | SAMN33548770 | Male | Mature | |

### Supporting Text 1: Extended laboratory and genome assembly methods

#### Long-read sequencing

High molecular weight (HMW) DNA was extracted from one female of each genotype (*Ao*, *Io*, *oo*) for morph-specific genome assemblies of *I. elegans* (Biosample accessions: SAMN33548774, SAMN33548775, SAMN33548776). Whole-thorax samples (15-20 mg) were submerged in liquid nitrogen and grounded into fine dust using a ceramic pestle. The ground samples were used as input for the Nanobind® Tissue Big Extraction Kit (Cat. No. NB-900-701-01, Circulomics Inc. (PacBio), MD, USA). We followed the Insect DNA Extraction Protocol v.0.20a, except for increased Proteinase K to 30 $\mu$l and increased incubation time to 1.5 h. DNA was thereafter treated with the Short Read Eliminator Kit (Cat. No. SS-100-101-01, Circulomics Inc. (PacBio), MD, USA), to progressively deplete fragments $< 25$ kb. DNA purity was measured from one 1 $\mu$l aliquote using a Nanodrop 800 (Thermo Fisher Scientific, MA, USA). DNA concentration was approximately quantified using three 1 $\mu$l aliqoutes and a Qubit 2.0 fluorometer (dsDNA BR; Invitrogen, CA, USA).

Sequencing libraries were constructed from each HMW DNA sample for the Nanopore LSK-110 ligation kit (Oxford Nanopore Technologies, UK). The manufacturer's protocol was modified as follows: 1) New England Biolabs (NEB) end preparation/DNA repair times were extended to 30 min at 20°C and 30 min at 65°C, 2) the elution of magnetic beads after end preparation/DNA repair was performed at 37°C and extended to ~40 min. DNA extractions, size selection, end-preparation and DNA repair were conducted at the Department of Zoology of Stockholm University. Adapter ligation and sequencing were carried out at the Uppsala Genome Centre (NGI), hosted by SciLife Lab. Each sample was sequenced on a PromethION R10.4

HMW isolation from homozygous *A* and *O*-like females of *I. senegalensis* (Biosample accessions: SAMN33548777 and SAMN33548778) were similar to the procedures described above, except for the following differences. Ground tissue samples were used as input for the Monarch® HMW DNA Extraction Kit for Tissue (Cat. No. T3060S, New England BioLabs Inc., MA, USA). Samples were incubated with lysis buffer and 20 $\mu$l of Proteinase K for 90 min, including 15 min of agitation at 1,400 rpm. DNA isolates were sheared using a Megaruptor® 3 (Cat. No. B06010003, Diagenoge Inc., NJ, USA), targeting a size of 90-100 kb (*O*-like sample) and 100-130 kb (*A* sample). Sample shearing, size selection, library preparation and sequencing were carried out by the Integrated Genomics Platform, Genome Institute of Singapore (GIS), A-STAR, Singapore. Each sample was sequenced on a PromethION R9.4 flow cell, with 2 nuclease washes and three library loadings.

#### Short-read sequencing

DNA isolation, library preparation, and sequencing were conducted in two batches. The first group included whole-thorax tissues from 19 *I* females and 16 *A* females, which were disrupted using a TissueLyser (Qiagen, Germany) or a mortar and pestle. DNA isolation followed the standard protocol of the QIAGEN Supplementary Protocol (*Purification of total DNA from insects using the DNeasy®Blood and Tissue Kit*) with the modifications described below. We increased Proteinase K to 40 $\mu$l and extended incubation time to three hours. After incubation, 130 $\mu$l of Buffer P3 (Cat. No. 19053, Qiagen, Germany) was added to the samples and after five minutes of incubation the samples were centrifuged for five minutes at 14,000 rpm. The resulting lysate was then transferred into a QIAshredder Mini spin column (Cat. No. 79656, Qiagen, Germany) and centrifuged at 14,000 rpm for two minutes. The lysate was then transferred into a new tube, adding 4 $\mu$l of RNase A (Cat. No. 19101, Qiagen, Germany), and incubating for two minutes. Lastly, we replaced the elution buffer provided with Buffer EB (Cat. No. 19086, Qiagen, Germany) and decreased the elution volume to 60 $\mu$l. DNA concentration was approximately quantified using a Nanodrop 2000 (Thermo Fisher Scientific, MA, USA), and degradation was inspected using 1% agarose gel scans with a lambda ladder (Cat. No. SM0102, Thermo Fisher Scientific, MA, USA).

The second batch included 19 whole-thorax samples from *O* females and three samples from *A* females. Tissues were disrupted using a pestle, in 200 $\mu$l of Lysis Buffer and 25 $\mu$l of Proteinase K, and incubated at 56 °C overnight. DNA was isolated using the KingFisher Cell and Tissue DNA Kit (Cat no. N11997, ThermoFisher Scientific) and the robotic Kingfisher Duo Prime purification system, following the manufacturer's protocols. DNA quality and quantity were assessed using a Nanodrop 800 (Thermo Fisher Scientific) and a Qubit 2.0 fluorometer (dsDNA BR; Invitrogen, Carlsbad, CA, USA).

Sequencing libraries for all samples were prepared from 2 $\mu$g of DNA, using the TrueSeq Nano DNA sample preparation kit (Cat. No. 20015964, Illumina Inc., CA, USA) and unique dual indexes (Cat. No. 20022370, Illumina Inc., CA, USA), targeting an insert size of 350 bp. The library preparation was performed according to the manufacturers' instructions. Libraries were sequenced on NovaSeq 6000 system with paired-end read length of 150 bp, using a S4 flowcell and v 1.5 sequencing chemistry. Library preparation and sequencing was conducted by SciLifeLab at the Uppsala Genome Centre ( NGI).

#### *I. senegalensis* DNA extraction and pooled sequencing

*I. senegalensis* DNA (biosample acessions: SAMN33548779, SAMN33548780) was extracted from muscle tissues in thoraxes using Maxwell® 16 LEV Plant DNA Kit (Cat. No. AS1420, Promega, WI, USA). For *A* and *O*-like, equal amounts of DNA from each of the 30 individuals was pooled. The pooled DNA was sequenced at the BGI Japan (Kobe, Japan) using Illmina HiSeq 4000 platform with 100 bp paired-end reads.

#### RNA extraction and sequencing

Whole-thorax samples were grounded into a fine powder using a TissueLyser. The samples were kept frozen at all times using liquid nitrogen and used as input for the Spectrum™ Plant Total RNA Kit (Cat. No. STRN50, Sigma Aldrich, MO, USA), including DNase I treatment (Cat. No. DNASE10, Sigma Aldrich, MO, USA). RNA concentration and quality were measured from 1 $\mu$l aliquot using a bioanalyzer (2100 Bioanalyzer System, Eukaryote Total RNA Nano Series II.xsy, Agilent Technologies, CA, USA). Library preparation and sequencing were performed by SciLife Lab at the Uppsala Genome Centre (NGI). Sequencing libraries were prepared from 300 ng of RNA, using the TrueSeq stranded mRNA library preparation kit (Cat. No. 20020595, Illumina Inc) including polyA selection and unique dual indexing (Cat. No. 20022371, Illumina Inc.), according to the manufacturer's protocol. Sequencing was performed on the Illumina NovaSeq 6000 SP flowcell with paired-end reads of 150 bp.

**Table S3.** Parameters used in Shasta assemblies of morph-specific genomes in *Ischnura elegans*. Four different configurations were tried for each assembly (T1-T4), which comprise alternative modifications of the configuration recommended by June 2020, for long-read Nanopore data. Non-applicable options and options left to the program default values in all configurations are omitted.

| Option | June2020 | T1 | T2 | T3 | T4 |
|---|---|---|---|---|---|
| Reads.minReadLength | 10000 | 10000 | 7000 | 7000 | 7000 |
| kmers.k | 14 | 14 | 14 | 14 | 14 |
| MinHash.minBucketSize | 5 | 4 | 5 | 5 | 4 |
| MinHash.maxBucketSize | 30 | 10 | 30 | 30 | 10 |
| MinHash.minFrequency | 5 | 2 | 5 | 5 | 2 |
| Align.maxSkip | 30 | 30 | 100 | 100 | 30 |
| Align.maxDrift | 30 | 30 | 100 | 100 | 30 |
| Align.maxTrim | 30 | 30 | 100 | 100 | 30 |
| Align.minAlignedMarkerCount | 400 | 100 | 10 | 10 | 100 |
| Align.minAlignedFraction | 0.55 | 0 | 0.1 | 0.1 | 0 |
| Align.downsamplingFactor | 0.05 | 0.1 | 0.05 | 0.05 | 0.1 |
| Align.sameChannelReadAlignment.suppressDeltaThreshold | 30 | 30 | 30 | 30 | 30 |
| ReadGraph.creationMethod | 0 | 0 | 2 | 2 | 0 |
| MarkerGraph.minCoverage | 10 | 10 | 0 | 0 | 10 |
| MarkerGraph.refineThreshold | 6 | 6 | 6 | 6 | 6 |
| MarkerGraph.crossEdgeCoverageThreshold | 3 | 3 | 3 | 3 | 3 |
| MarkerGraph.simplifyMaxLength | 10,100,1000,10000,100000 | 10,100,1000,10000,100000 | 10,100,1000,10000,100000 | 10,100,1000,10000,100000 | 10,100,1000,10000,100000 |
| Assembly.detangleMethod | 1 | 1 | 2 | 1 | 1 |
| Assembly.consensusCaller | Bayesian:guppy-3.6.0-a | Bayesian:guppy-3.6.0-a | Bayesian:guppy-3.6.0-a | Bayesian:guppy-3.6.0-a | Bayesian:guppy-3.6.0-a |

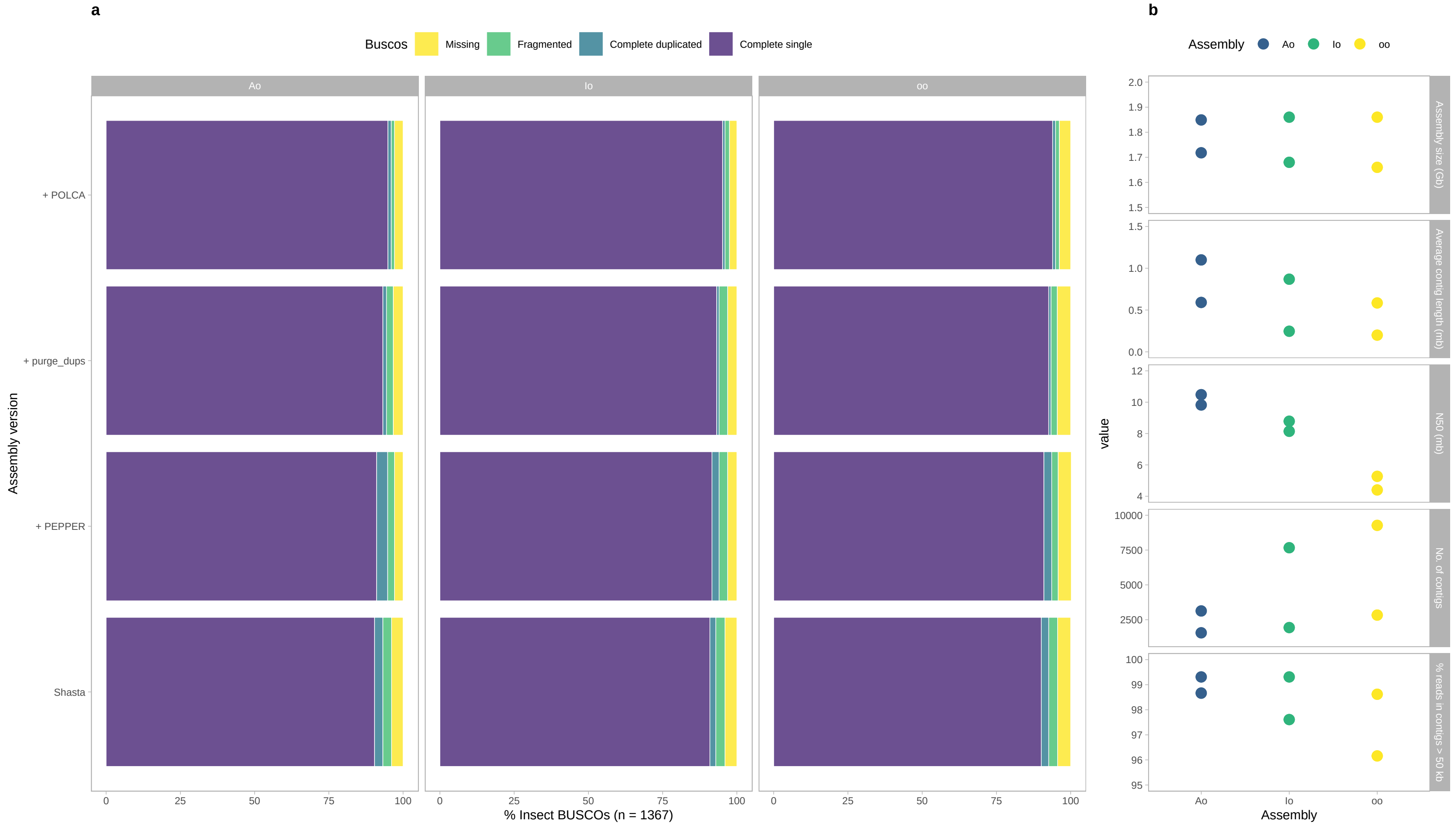

**Figure S1.** Quality assessment of morph-specific assemblies of *Ischnura elegans* at different stages of polishing. **a** Completeness of conserved insect genes. **b** Assembly statistics reflecting total size (in Gb) and contiguity.

**a**

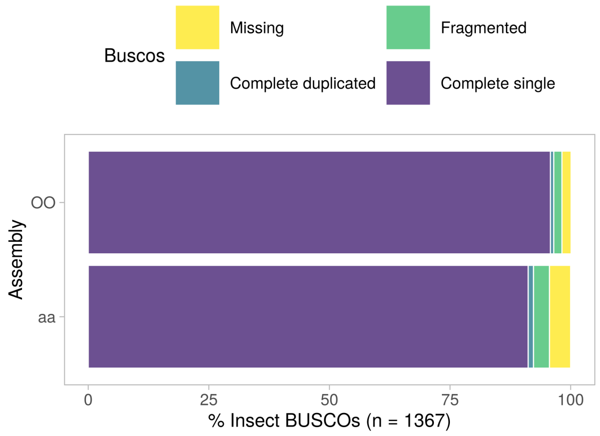

**b**

| Statistic | aa Assembly | OO Assembly |
|---|---|---|
| N 50 (mb) | 0.71 | 4.51 |
| Average contig length (mb) | 0.23 | 1.12 |
| No. of contigs | 6550 | 1409 |
| % reads in contigs>50 kb | 96.15 | 99.49 |
| Assembly size (Gb) | 1.53 | 1.58 |

**Figure S2.** Quality assessment of morph-specific assemblies of *Ischnura senegalensis*. **a** Completeness of conserved insect genes. **b** Assembly statistics reflecting total size (in Gb) and contiguity.

### Supporting Text 2: Genotyping the DToL reference assembly

The genotype of the specimen on which the DToL reference assembly is based was unknown upon data collection. To determine the genotype of this individual, we started by mappping its raw long-read data to our *A* assembly and quantified read-depth as for our own data. The distribution of standardized read depth values across the unlocalized scaffold 2 of chromosome 13 in our *A* assembly indicated that the DToL sample included at least one copy, and presumably only one copy, of the *A* allele (Fig. S3a).

To investigate which allele was preserved in the primary DToL assembly, we then aligned this assembly to our *A* morph assembly. The alignment along the unlocalized scaffold of chromosome 13 was visibly broken in the two regions of genomic content we previously identified as present in *A* samples, but absent in *O* samples (Fig. S3b). This indicated that while the *A* allele is present in the raw data, it may have been purged from the final haploid assembly. We therefore also aligned our *A* assembly to the haplotigs filtered from the main DToL assembly, and found a single haplotig (RAPID_106) with a nearly continuous alignment between the two (Fig. S3b). Moreover, this DToL haplotig contained most of the significant *k*-mers identified as *A*-unique content in the *k*-mer based GWAS (Extended Data Table 1), whereas the *A*-specific and *A*-and-*I*-specific *k*-mers were nearly absent from the primary DToL assembly (Extended Data Table 1). We thus concluded that the primary DToL assembly captured the *O* allele, whereas the *A* allele, also present in the DToL specimen, is represented in the alternative haplotig RAPID_106.

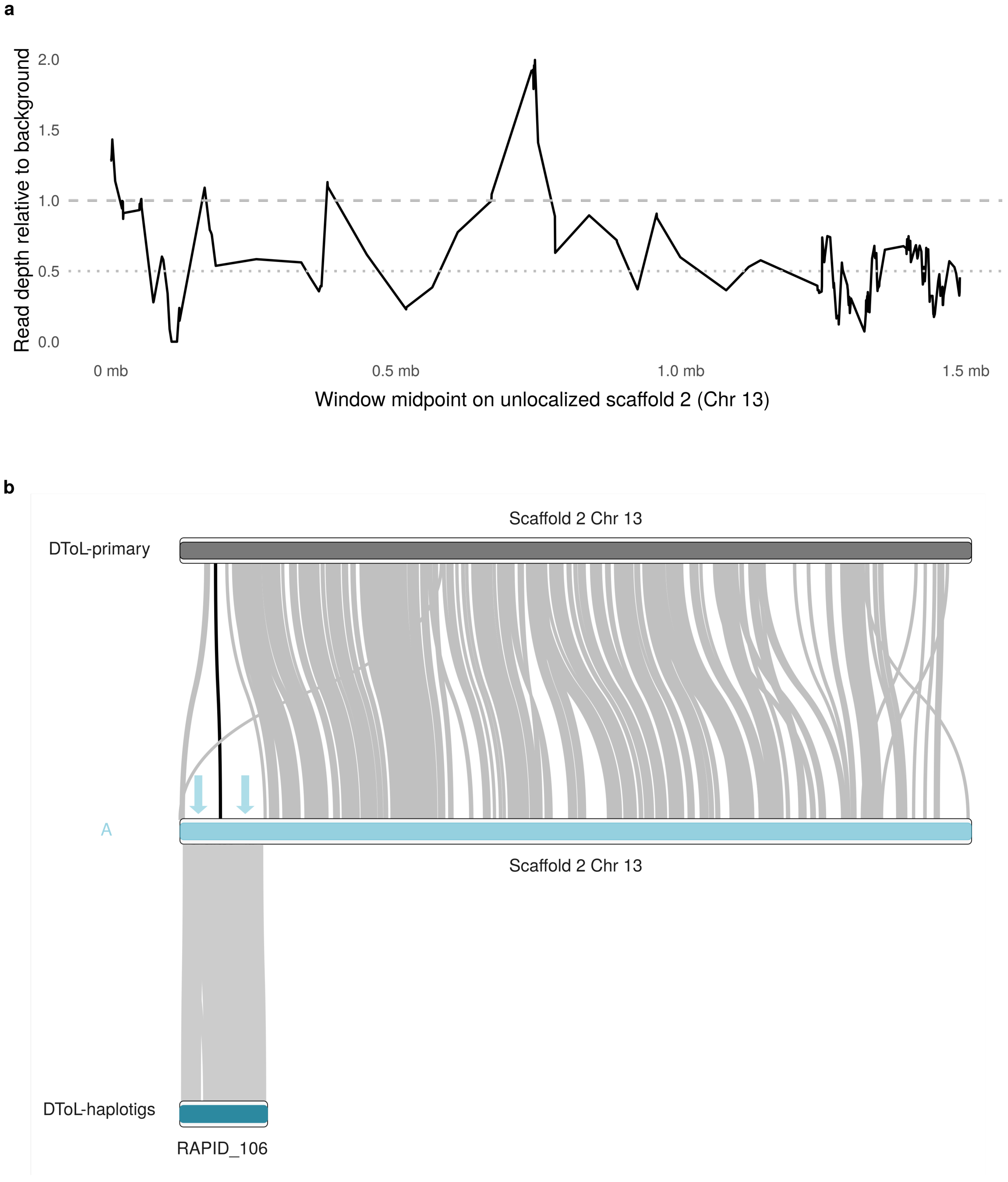

**Figure S3.** Genotyping the DToL reference assembly. **a** Standardized read depth of the raw Darwin Tree of Life (DToL) data along the first 1.5 mb of the unlocalized scaffold 2 of chromosome 13 in our *A* morph assembly. Read depth is standardized relative to background coverage. The dotted and dashed gray lines show read depths of 0.5 and 1.0, respectively. **b** Alignment between the unlocalized scaffold 2 of chromosome 13 in our *A* morph assembly (middle) and the Darwin Tree of Life (DToL) assembly. The top bar corresponds to the unlocalized scaffold 2 of chromosome 13 in the primary DToL assembly and the bottom bar shows the alternative haplotig RAPID 106. The windows identified as genomic content present in *A* females but absent in *O* are marked with arrows.

### Supporting Text 3: Potential recombinants or incomplete penetrance

Two samples (one identified as *A* and one identified as *O*) contained genomic content and SV signatures that were associated with an alternative female morph. These samples could reflect phenotyping errors, which are unlikely. Alternatively, these specimens may indicate instances of recombination within the morph locus, although such recombination should be rare due to the strong linkage disequilibrium across of chromosome 13 (Extended Data Fig. 5). Finally, individuals with mismatched phenotypes may be the product of incomplete penetrance of morph alleles. Incomplete penetrance has been recorded in sex-determination genes through their interactions with genes downstream in the sex determination cascade[1], and it is suspected that epistatic interactions might also be responsible for incomplete penetrance of the colour polymorphism controlled by an autosomal supergene in the land snail *Cepaea nemoralis*[2]. If allelic differences in the morph SVs indeed influence penetrance, these effects could be leveraged in future studies to narrow down the genetic basis of morph determination.

We further examined these potentially outliers using PCA analyses of allele frequencies at the morph loci. With these analyses, we visualized how short-read samples from different morphs distribute along two main axes of SNP variation, and whether the outliers seem to be intermediate between two morphs or group within samples of a mismatched morph (Fig. S4). We conducted PCA analyses based on the unlocalized scaffold 2 of chromosome 13 of both the *A* reference assembly (1 bp - 1.6 mb) and the *I* assembly (3.5 - 3.7 mb). In each case, we used a vcf file filtered with the same criteria as for the GWAS analyses. We performed linkage pruning on these files in *PLINK* v 1.9[3], using a window size of 50 kb a step size of 10 bp, and setting the $R^2$ threshold to 0.1. We then conducted the PCA analyses also using *PLINK* v 1.9[3] and plotted sample values on the first two principal components using the *ggplot2* v 3.4.1[4] package in *R* v 4.2.2[5]. Both outlier samples seem to cluster with a mismatched morph, particularly when using the *I*-morph assembly as mapping reference (Fig. S4), and thus point at a potential role of incomplete penetrance that warrants further study.

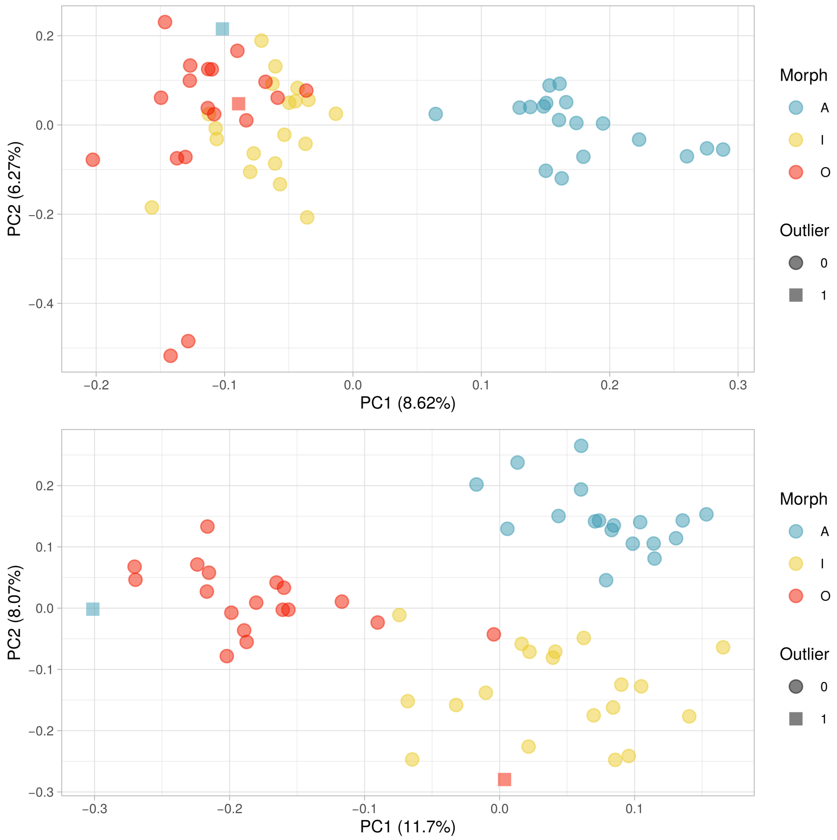

**Figure S4.** PCA analysis of allele frequencies on **a** the morph locus of the *A*-morph assembly and **b** the morph locus (differentiating *A* and *I* from *O*) of the *I*-morph assembly. 57 short-read genomic samples visually identified to morph are included. Two samples, labelled as outliers, exhibited genomic content and SV signatures characteristic of a different morph.

#### Supporting Text 4: Structural variation between morphs

In order to investigate how the distinct morph haplotypes may have formed, we screened SVs in short-read samples from *A* and *I* females against the *O* genome assembly. Here, we shifted to the *O* assembly, as mapping reference because comparative analysis indicated this is the ancestral morph in the genus *Ischnura*[6]. We also focused on the start of the unlocalized scaffold 2 of chromosome 13, where the original divergence between *O* and *A* is presumed to have taken place. This hypothesis is based on the findings that both the *A* and *I* haplotypes of *I. elegans* differ from the *O* allele in this region (Fig. 2b-c), and that the *A* alleles in both *I. elegans* and *I. senegalensis* differ from the *O* and *O-like* alleles respectively along the same region (Fig. 5b-c). We uncovered evidence of a ~20 kb inversion, shared by *A* and *I* females, but absent in all but one sample phenotyped as *O* (Extended Data Fig. 2). Importantly, this finding is robust to whether SVs are called against our own *O* assembly, or the DToL reference genome, which also captures the *O* allele in its primary assembly (Extended Data Fig. 2).

We also noted a spike in read depth coverage, specially in *A* samples, at the breakpoints of the putative inversion (Extended Data Fig. 2), suggesting that part of this sequence may have propagated to multiple locations in the *A* genome. To examine this possibility, we extracted all reads mapping to the breakpoints and reasoned that break-point reads derived from *A* and *I* samples should map to multiple positions on the *A* assembly. As suggested by read depth coverage, the break-point reads derived from *A* samples mapped to three distinct regions in the *A* genome (Extended Data Fig. 3), whereas break-point reads derived from *I* samples mapped to fewer locations in the *A* assembly, most of them across between 50 kb and 150 kb (Extended Data Fig. 3). Read depth coverage and mapping location of break-point reads thus implied that a relatively short sequence, probably a TE spanning the inversion break-point, has proliferated in the *A* genome.

While SV calling across the entire morph locus was likely precluded by its extension[7] (~1.5 mb in *A* and ~3.5 mb in *I*), we uncovered one morph-specific SV within the locus, that was supported by both short and long-read data. The genomic region shared by all morphs (between ~0.6-1.0 mb in the *A* haplotype) contains a ~6.7 kb signature of both duplication and deletion in *I* and *O* samples against the *A* reference (Fig. S5). Such an occurrence of both duplication and deletion signatures in the same region is consistent with an event of non-homologous recombination of a sequence repeated in tandem, suggesting additional recombination along repetitive regions may have shaped morph divergence in *Ischnura*.

Next, we examined the evidence for a translocation from *A* to *I*, as suggested by the assembly alignment and *k*-mer mapping (Fig. 3b; 4a). To confirm that this is not a product of an error in the assembly of the

*I* genome, we called SVs using the raw long-read data from a *Io* individual against the *A* assembly. Here, we expected a signature of an inversion between the original position in *A* females and the translocated position in *I* females. Because the morph assemblies differ in length, the ending position of the putative translocation in the *I* assembly (~3.76 mb) aligns to the *A* assembly at ~5.54 mb. Our analysis supported an inversion comprising this whole ~5.54 mb region (Extended Data Fig. 4), which is consistent with an inverted translocation, a translocation of the original inversion or an inversion of 5.54 mb. As there is no evidence of morph divergence beyond ~1.5 mb on the *A* assembly (Fig. 2-3), the SV call is more appropriately interpreted as evidence of either translocation scenario (see also Fig. S6). Translocation events can be facilitated by TE activity, which propagates homologous sequences in different genomic locations and can thus result in ectopic recombination[8,9]. Notably, chromosome 13 of the *I. elegans* genome is enriched for TE sequences (Extended Data Fig. 4), athough this pattern varies between TE families (Fig. S7).

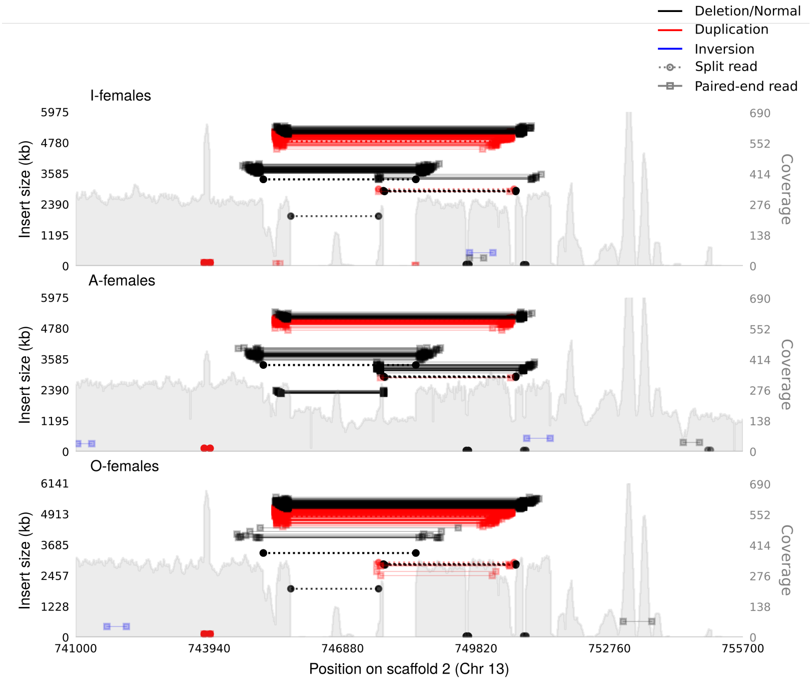

**Figure S5.** Read mapping and sample coverage showing signatures of both duplication and deletion in *I* and *O* samples against the *A* assembly, in the central region of the morph locus (0.6-1.0 mb) that is shared by all female morphs. As most of the *A* samples come from heterozygous individuals the signatures of both duplication and deletion are present in these samples, but are supported by fewer reads compared to *I* and *O* samples.

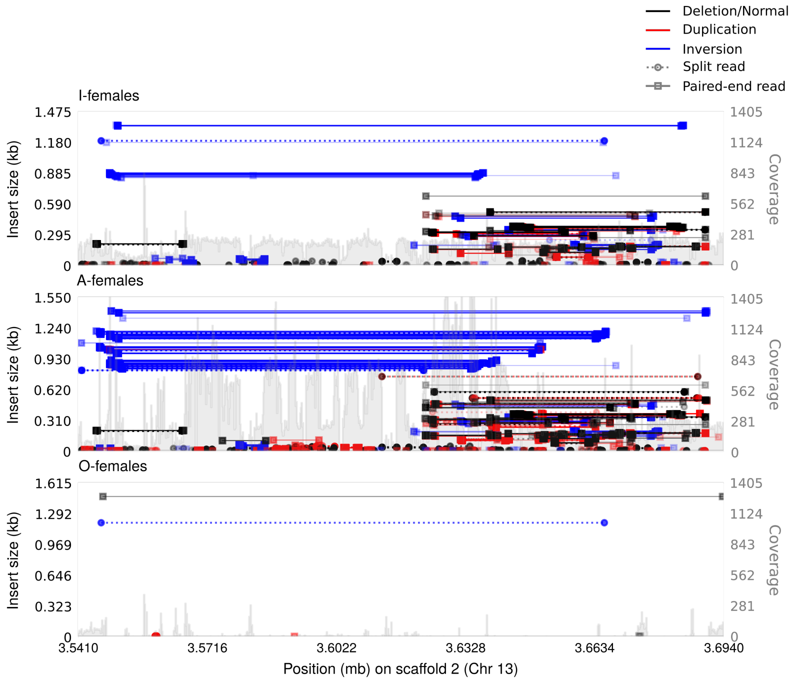

**Figure S6.** Read mapping and sample coverage at the putative translocation site on scaffold 2 of chromosome 13 of the *I* assembly. Mapped reads from *A* samples show multiple inversion signatures, whereas *I* samples have fewer signatures. A single *O* sample exhibited a pattern similar to *I* females, but was excluded here for clarity (see Discussion).

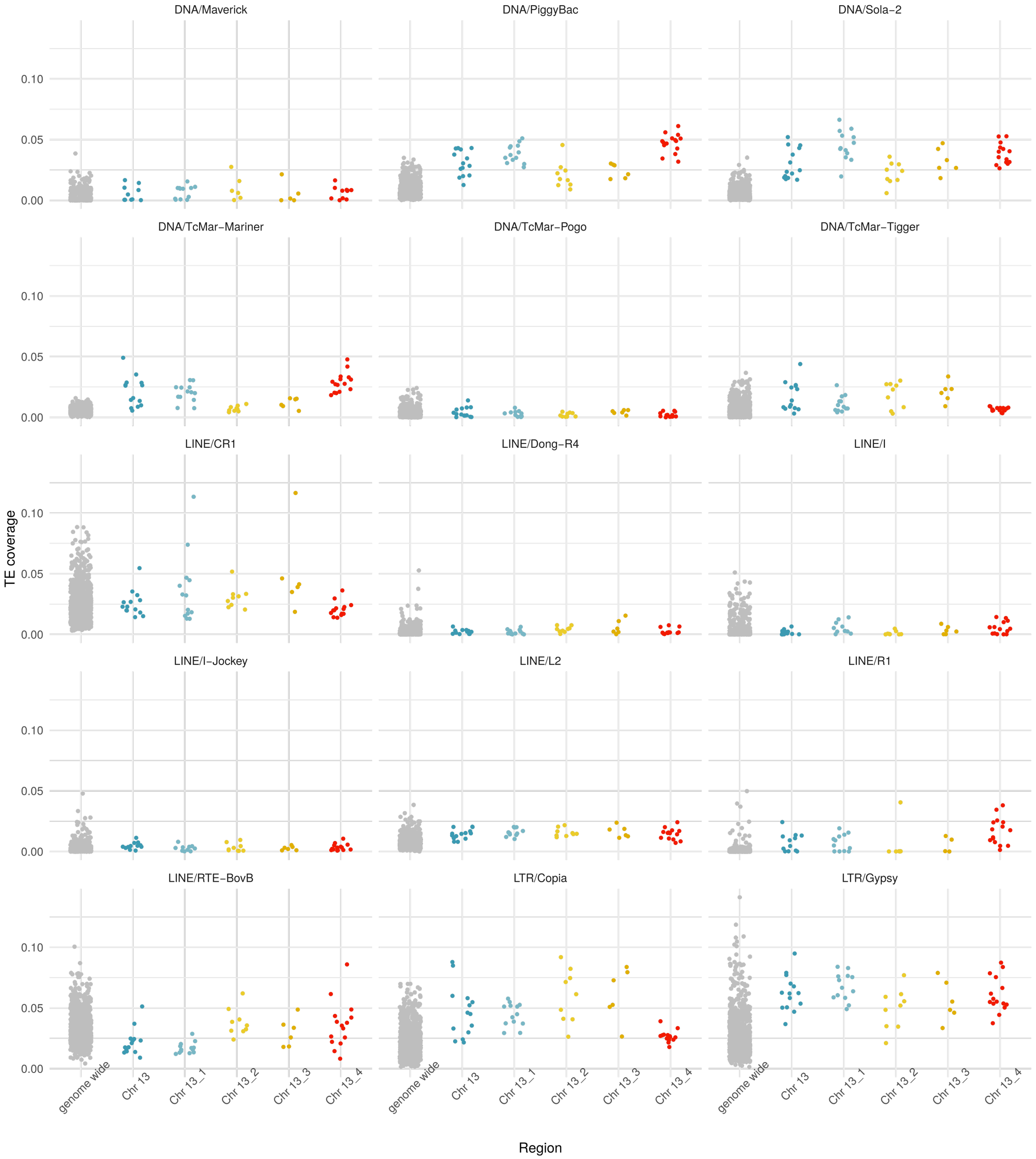

**Figure S7.** Proportion of the *A* assembly covered by the most abundant TE families. Coverage is calculated in 1.5 mb regions for the main assembly of chromosome 13, the four unlocalized scaffolds of chromosome 13 and the rest of the genome.

#### Supporting Text 5: A shared genomic basis of inter-sexual mimicry

In the main text, we report a difference in the standardized number of mapped reads (read depths) between *A* and *O*-like sequencing pools of *I. senegalensis* along the morph locus of *I. elegans* (Fig. 5b). These results show higher coverage in the *I. senegalensis A* pool of regions of genomic content unique to *A* females of *I. elengas*. To confirm that this difference in read depth between female morphs in *I. senegalensis* is specific to the morph locus, we compared it to the distribution of read-depth differences in the rest of the genome. We found that the *A* pool of *I. senegalensis* has relatively high mapping coverage along the morph locus, compared to the genome-wide pattern (Fig. S8).

We also mapped the data from both *I. senegalensis* pools to the *I. elegans O* assembly, and used Samplot[10] to inspect the region where *A* and *I* females of *I. elegans* carry a ~20 kb inversion signature and elevated read-depth coverage compared to the *O* morph (Extended Data Fig. 2). We identified a signature of the same inversion and elevated read-depth in the *A* pool of *I. senegalensis* (Extended Data Fig. 7). As in *A* females of *I. elegans*, the breakpoint reads from the *A* pool mapped to multiple locations across the *A* assembly (compare Extended Data Fig. 3 to Extended Data Fig. 7b). Together, these results imply that an inversion and propagation of a repetitive sequence in chromosome 13 is shared by *A* females of both *I. elegans* and *I. senegalensis*.

Finally, we generated and aligned *de novo* assemblies for two homozygous females of *I. senegalensis* in Singapore (Extended Data Fig. 1). Female-morph dominance is reversed in *I. senegalensis*, where the *O* allele is dominant over *A*[11]. Therefore, all females with the *A* phenotype are homozygous for the *A* allele. To obtain a homozygous *O*-like genome, we designed primers for an *A*-specific sequence shared by *I. elegans* and *I. senegalensis* (forward: CGCGGTATGATATGGTCCGA, reverse: GGCTGCTTACACCAATGCAA). For clarity, we plot only one contig of each *I. senegalensis* assembly for regions of the *I. elegans* genome that have multiple alignments, choosing contigs based on the total alignment length. These alignments show that the *A*-unique genomic region in *I. elegans* is partly shared by *A* females but not *O*-like females of *I. senegalensis* (Fig. 5c).

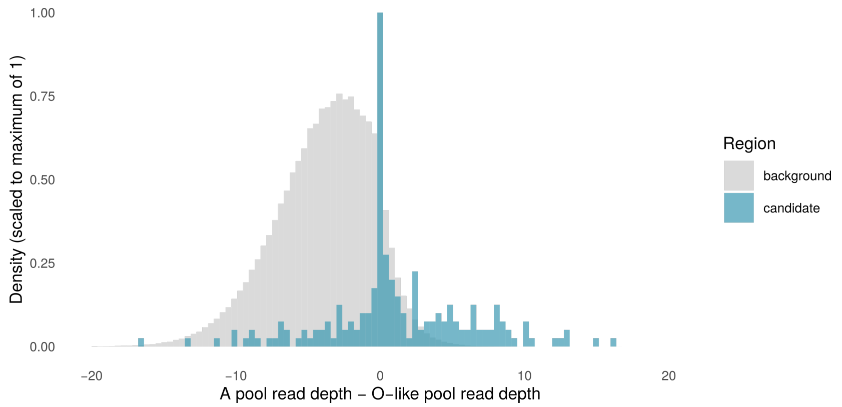

**Figure S8.** Supporting evidence for a shared molecular basis of inter-sexual mimicry in *I. elegans* and *I. senegalensis*. Difference in read-depths between the *A* pool and the *O*-like pool of *I. senengalensis*, across 500 bp windows on the *I. elegans* *A*-morph assembly.

#### Supporting Text 6: Predicted genes in the morph locus

As male-mimicry is characterized by unique genomic content in the two species (Fig. 3; 5), we first investigated whether any genes are present and expressed in *A* females of both *I. elegans* and *I. senegalensis*, but not in *O* and *O*-like females. Seven gene models are encoded in the ~1.5 mb morph locus (of the *A* assembly) and expressed in both species (Fig. 6a). Of these, four are located in a genomic region absent in *O* females and are either unique to *A* or shared by both *A* and *I* females (Fig. 6a). Consequently, transcripts derived from these genes were either absent from the *O*-morph DToL transcriptome, or mapped to transcripts located in other chromosomes or scaffolds (Table S4). In *I. elegans*, all of these transcripts were expressed exclusively in *A* females or in *A* and *I* females, if present in the two morphs (Fig. 6b; S9; Table S4). However, two *A*-associated genes in *I. elegans* were expressed in both *A* and *O*-like morphs of *I. senegalensis* (Fig. 6b; S10), suggesting that these gene models are unlikely causative of the sexual-mimicking phenotype shared by the two species. The *A*-specific expression and mapping of these genes in *I. elegans* may be a consequence of translocation to the *A* haplotype after the split between the ancestors of the two species more than 5 Ma[6].

We noted that a gene uniquely present in *A* and *I* females (Afem.4094) and two other genes outside the morph locus undergo expression changes in *I* females that recapitulate adult colour changes, such that the gene's expression pattern is initially more similar to *A* females and becomes more *O*-like with sexual maturation. Afem.4094 is similarly expressed in immature *A* and *I* females, but ceases to be expressed in mature *I* females (Fig. S11). Thus, upon sexual maturation, *I* females resemble *O* females, in which this gene is absent and never expressed. The two other genes code for THEM6 (Thioesterase family member 6) and an ELOVL (elongation of very long chain fatty acid) protein, both likely involved in fatty acid metabolism[12–14] (Table S5). These genes are downregulated in adult development of both *I* and *O* females, while remaining highly expressed in mature *A* females and males. The synthesis of structural waxes, which consist of long-chain fatty acids, is responsible for blue coloration in other species of Odonata[15,16], and may thus potentially contribute to the loss of blue colouration with adult development in *I* females.

Next, we examined the three gene models that are shared between the *A* and *O* haplotypes in both species (Fig. 6a). One of these gene models (Afem.4119 in our *A* assembly, LOC124172681 in the reference assembly), annotated as a zinc finger protein 665-like, was expressed exclusively in males of both species (Fig. 6b, Fig. S9 -S10). The other two genes models (Afem.4093 and Afem.4111) also encode zinc-finger domain proteins (Fig. 6b; Table S4). These two genes are expressed in females, but there was no evidence of differential expression between morphs in the adult stage (Fig. S9-S10; Table S6). These genes may be nonetheless important for the evolution and function of the morph locus.

The gene Afem.4093 flanks the genomic region uniquely shared by *A* and *I* females, and is highly similar (95% identity) to a second gene model located ~3.5 mb downstream on the same scaffold (Fig. S12). We found no evidence of differential expression between morphs in either of these flanking genes (Fig. S13). Nonetheless, in both the *A* and *I* haplotypes a similar coding gene and Jockey element flank the region of novel genomic content shared by these derived morphs (Fig. 6a for *A*; Fig. S12 for *I*). Even though these regions occupy different positions on the chromosome, their sequence similarity may have facilitated an ectopic recombination event, leading to the origin of *I* females.

The gene model Afem.4111 (LOC124172682 in the reference assembly) is located in the island of shared genomic content bounded by SVs in our *A*-morph assembly (Fig. 4a; 6a). Functional annotation of insect orthologues indicates that Afem.4111 is likely a DNA-binding transcription factor, that regulates the expression of mRNA. We assembled two isoforms for this gene. The long isoform (Afem.4111.2) contains all exons, and encodes a AD-type zinc finger domain, an uncharacterized domain, and a series of C2H2 zinc fingers (Fig. S14a). In the short isoform (Afem.4111.1), only some C2H2 zinc-finger coding exons are retained.

We used *OrthoFinder* v 2.5.2[17] to identify orthologues of Afem.4111 in other insect proteomes and in other chromosomal locations in the *I. elegans* genome. For this analysis, we used the proteome of another species in the order Odonata (*Ladona fulva*) and representative proteomes of hemimetabolous (*Acyrthosiphon pisum*, *Frankliniella occidentalis*, and *Zootermopsis nevadensis*) and holometabolous (*Bicyclus anynana*, *Tribolium castaneum*, and *Drosophila melanogaster*) insects. Orthologue group inference suggested that Afem.4111 is part of a gene clade particularly diversified in Odonata (dragonflies and damselflies) with most paralogues clustered in chromosome 13 (Fig. S14b).

Finally, we used genotype plots to identify fixed SNPs between morphs in the three protein coding genes that are situated in the morph locus and are shared between morph haplotypes (Afem.4093, Afem.4111 and Afem.4119). Genotypes of short-read samples at the coding regions of these genes were based on the same vcf file as for the GWAS analyses, an plotted using the *GenotypePlot*[18] package in *R*. Afem.4111 was the only of the shared gene models with non-synonymous SNPs between morphs, in all cases differentiating *A* from both *I* and *O* females (Fig. S14a). However, all of these non-synonymous mutations were accumulated in the uncharacterized regions of the gene (Fig. S14a), and none were shared between *A* females in the two species.

**Table S4.** Annotation of transcripts and inferred coding sequences of genes in the morph locus of *A* females and expressed in adults of both *I. elegans* and *I. senegalensis*. We blasted transcripts in the morph locus of *A* females against the DToL reference assembly and determined whether genes were syntenic in the two haplotypes, uniquely present in the *A* haplotype or unique to the *A* haplotype but having some similarity to genes elsewhere in the DToL reference. Coding sequences were inferred for all but one gene model (Afem.4099) in the morph locus, and translated into peptides. We report the best Swissprot hit for these peptides in insects, inferred functional domains, and functional annotations of the best EggNOG hit. For genes that are present and syntenic in both our *A* assembly and the DToL reference assembly, we report annotations based on the reference coding sequences, as these are based on a wider breadth of RNAseq data than our inferred peptides[19].

| Gene model | DToL reference | Swissprot hits | Functional domains | eggNOG annotation |
|---|---:|---:|---:|---:|
| Afem.4093 | syntenic to uncharacterized protein (LOC124172704) | No hits | Zinc finger, AD-type | Function unknown: Zinc-finger associated domain (zf-AD) (ENOG50392KJ) |
| Afem.4094 | absent, except for one exon sharing 245 bp with putative nuclease in chr 13 | RNA-directed DNA polymerase from mobile element jockey | Reverse transcriptase domain | Replication, recombination and repair: MAP kinase activity (ENOG5039XVS), Cell wall/membrane/envelope biogenesis:reverse transcriptase (ENOG5039VXI) |
| Afem.4099 | absent, except for one exon sharing 703 bp with ncRNA in chr 9 | NA | NA | NA |

| | | | | |
|---|---|---|---|---|
| Afem.4100 | 1291 bp shared with uncharacterized transcript in chr 3 | No hits | No annotated domains | Function unknown: Not available (ENOG502SFX0) |
| Afem.4103 | 223 bp shared with uncharacterized ncRNA in chr 6 | Retrovirus-related Pol polyprotein from type-2 retrotransposable element R2DM | Reverse transcriptase domain | Function unknown: Ribonuclease H protein (ENOG5039MXZ) Function unknown: Endonuclease-reverse transcriptase (ENOG503A2E1) |
| Afem.4111 | syntenic to gastrula zinc finger protein XlCGF57.1-like (LOC124172682) | Protein suppressor of hairy wing | Zinc finger, AD-type and Zinc finger, C2H2-type | Transcription: DNA binding (ENOG5038CKT) |
| Afem.4119 | syntenic to zinc finger protein 665-like (LOC124172681) | Krueppel homolog 1 | Zinc finger, AD-type | Transcription: Recombination hotspot binding (ENOG5038HRH) |

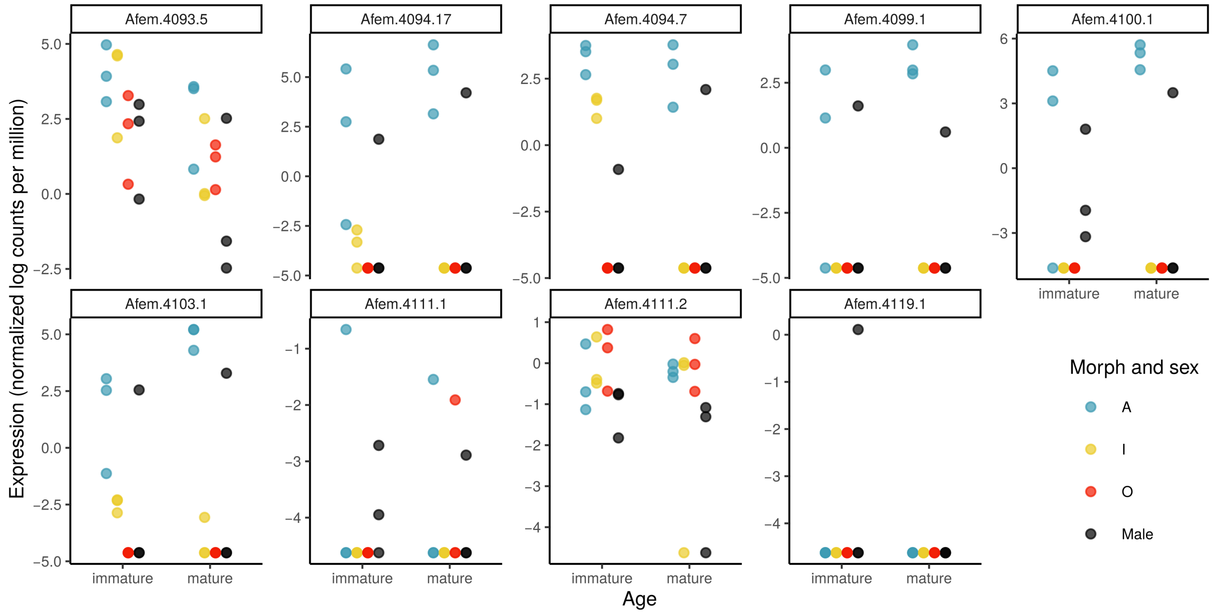

**Figure S9.** Expression of transcripts encoded in the morph locus of *I. elegans* and shared by *I. senegalensis*. Expression levels were quantified as log counts per million and normalized across whole-body samples of males and each of three female morphs.

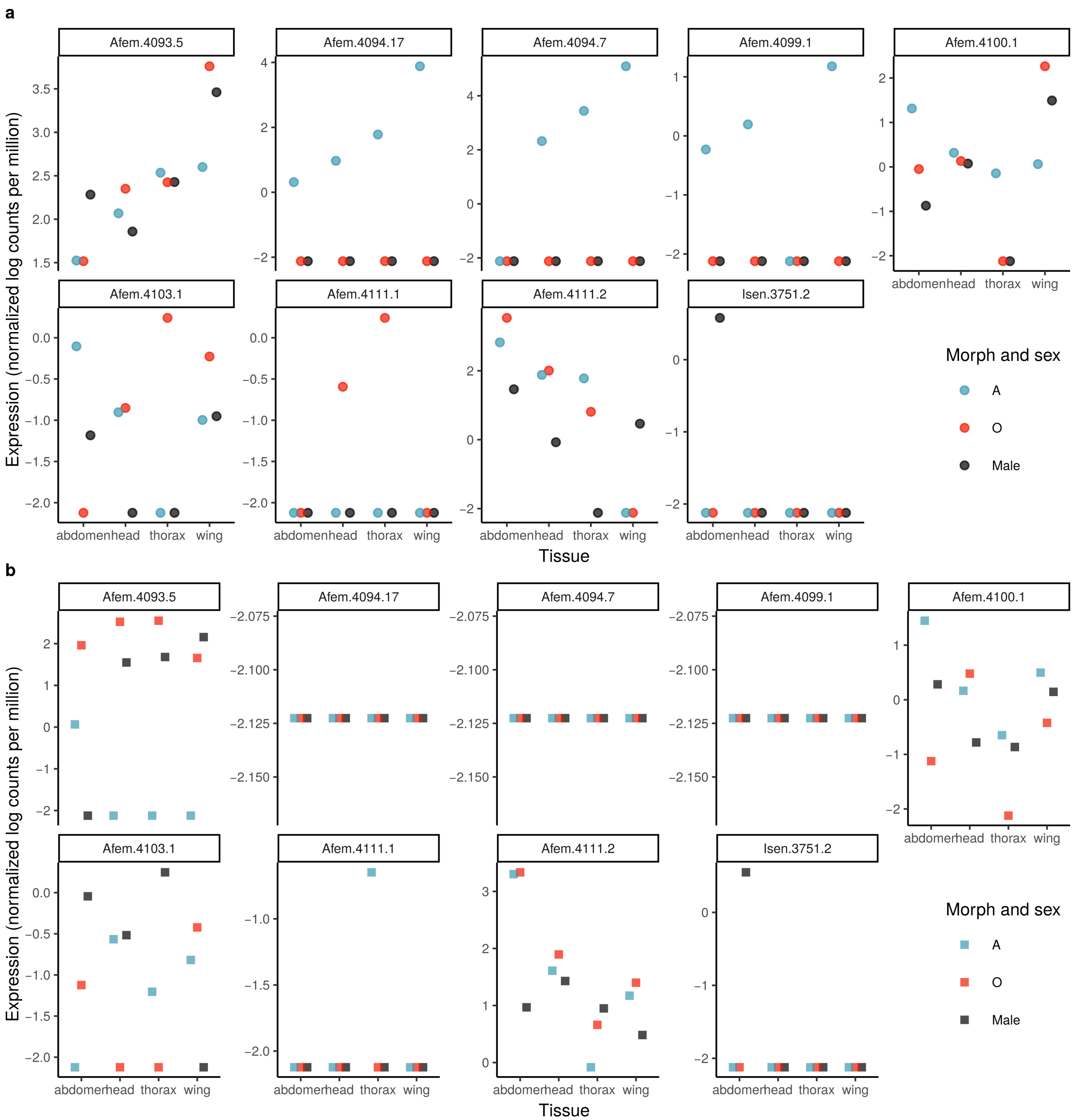

**Figure S10.** Expression of transcripts encoded in the morph locus of *I. elegans* in adult samples of *I. senegalensis*. Expression levels were quantified as log counts per million and normalized across samples of males and both female morphs. RNAseq data comes from a previous study (Okude et al. 2022) that included **a** one adult sampled upon emergence and **b** one adult sampled two days after emergence for each group. Transcript Isen.3751.2 is encoded within the gene model Afem.4119 assembled from RNAseq of *I. elegans*.

**Table S5.** Annotation of transcripts and inferred coding sequences of genes with expression changes in *I* females of *I. elegans*, reflecting their immature colour similarity to *A* females and their mature colour similarity to *O* females. We blasted transcript sequences, assembled using the *A* genome against the DToL reference assembly. Coding sequences were inferred and translated into peptides. We report the best Swissprot hit for these peptides in insects, inferred functional domains, and functional annotations of the best EggNOG hit.

| Gene model | DToL reference | Swissprot hits | Functional domains | eggNOG annotation |
| --- | --- | --- | --- | --- |
| Afem.4094 | absent, except for one exon sharing 245 bp with putative nuclease in chr 13 | RNA-directed DNA polymerase from mobile element jockey | Reverse transcriptase domain | Replication, recombination and repair: MAP kinase activity (ENOG5039XVS), Cell wall/membrane/envelope biogenesis:reverse transcriptase (ENOG5039VXI) |
| Afem.792 | Ischnura elegans elongation of very long chain fatty acids protein-like | Elongation of very long chain fatty acids protein 7 | ELO family | Lipid transport and metabolism: fatty acid elongation, saturated fatty acid (KOG3071) |
| Afem.13726 | THEM6 | THEM6 | HotDog domain of thioesterases | Function unknown: Thioesterase-like superfamily (KOG4366) |

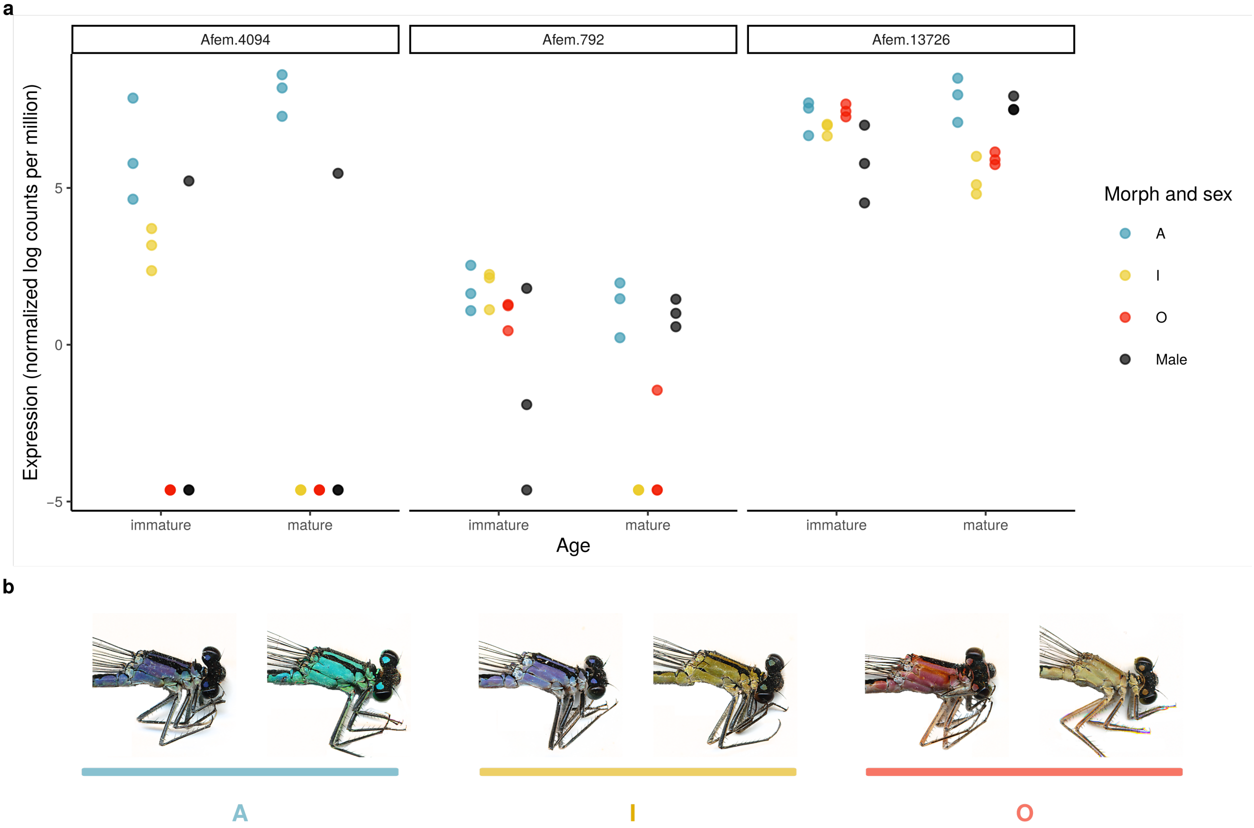

**Figure S11.** Three genes show expression changes in *I* females of *I. elegans* reflecting their immature colour similarity to *A* females and their mature colour similarity to *O* females. **a** Expression levels quantified as log counts per million, and normalized across whole-body samples of males and each of three female morphs. **b** Thoracic colouration of immature (left) and mature (right) females of each morph. A bright blue patch on the eighth abdominal segment is present in *A* females throughout adult development, but becomes obscure with sexual maturation in *O* and *I* females.

**Table S6.** DGE analysis of transcripts in the morph locus of *I. elegans*. Transcriptomic data were mapped to the *A*-morph assembly. P-values were computed from two-tailed exact tests and adjusted for multiple comparisons using the Benjamini and Hochberg's false discovery rate.

| Transcript | Stage | Contrast | log fold change | P-value |
|---|---|---|---|---|
| Afem.4093.5 | immature | A vs O | 1.786 | 0.153 |
| Afem.4093.5 | immature | A vs I | 0.075 | 0.951 |
| Afem.4093.5 | immature | I vs O | 1.711 | 0.170 |
| Afem.4093.5 | immature | A vs male | 1.956 | 0.119 |
| Afem.4093.5 | immature | I vs male | 1.880 | 0.133 |
| Afem.4093.5 | immature | O vs male | 0.170 | 0.889 |
| Afem.4093.5 | mature | A vs O | 1.956 | 0.120 |
| Afem.4093.5 | mature | A vs I | 1.731 | 0.166 |
| Afem.4093.5 | mature | I vs O | 0.225 | 0.853 |
| Afem.4093.5 | mature | A vs male | 2.049 | 0.104 |
| Afem.4093.5 | mature | I vs male | 0.319 | 0.793 |
| Afem.4093.5 | mature | O vs male | 0.093 | 0.939 |
| Afem.4094.17 | immature | A vs O | 12.687 | 0.000 |
| Afem.4094.17 | immature | A vs I | 8.136 | 0.008 |
| Afem.4094.17 | immature | I vs O | 4.551 | 0.062 |
| Afem.4094.17 | immature | A vs male | 3.788 | 0.134 |
| Afem.4094.17 | immature | I vs male | -4.348 | 0.094 |
| Afem.4094.17 | immature | O vs male | -8.899 | 0.004 |
| Afem.4094.17 | mature | A vs O | 14.270 | 0.000 |
| Afem.4094.17 | mature | A vs I | 14.270 | 0.000 |
| Afem.4094.17 | mature | I vs O | 0.000 | 1.000 |
| Afem.4094.17 | mature | A vs male | 3.044 | 0.213 |
| Afem.4094.17 | mature | I vs male | -11.226 | 0.001 |
| Afem.4094.17 | mature | O vs male | -11.226 | 0.001 |
| Afem.4094.7 | immature | A vs O | 11.994 | 0.000 |
| Afem.4094.7 | immature | A vs I | 1.891 | 0.383 |
| Afem.4094.7 | immature | I vs O | 10.104 | 0.001 |
| Afem.4094.7 | immature | A vs male | 5.959 | 0.022 |

*(continued)*

| Transcript | Stage | Contrast | log fold change | P-value |
|---|---|---|---|---|
| Afem.4094.7 | immature | I vs male | 4.068 | 0.087 |
| Afem.4094.7 | immature | O vs male | -6.035 | 0.018 |
| Afem.4094.7 | mature | A vs O | 11.647 | 0.000 |
| Afem.4094.7 | mature | A vs I | 11.647 | 0.000 |
| Afem.4094.7 | mature | I vs O | 0.000 | 1.000 |
| Afem.4094.7 | mature | A vs male | 2.544 | 0.252 |
| Afem.4094.7 | mature | I vs male | -9.104 | 0.002 |
| Afem.4094.7 | mature | O vs male | -9.104 | 0.003 |
| Afem.4099.1 | immature | A vs O | 10.393 | 0.009 |
| Afem.4099.1 | immature | A vs I | 10.393 | 0.009 |
| Afem.4099.1 | immature | I vs O | 0.000 | 1.000 |
| Afem.4099.1 | immature | A vs male | 1.757 | 0.545 |
| Afem.4099.1 | immature | I vs male | -8.636 | 0.018 |
| Afem.4099.1 | immature | O vs male | -8.636 | 0.018 |
| Afem.4099.1 | mature | A vs O | 11.973 | 0.005 |
| Afem.4099.1 | mature | A vs I | 11.973 | 0.005 |
| Afem.4099.1 | mature | I vs O | 0.000 | 1.000 |
| Afem.4099.1 | mature | A vs male | 4.371 | 0.178 |
| Afem.4099.1 | mature | I vs male | -7.602 | 0.030 |
| Afem.4099.1 | mature | O vs male | -7.602 | 0.032 |
| Afem.4100.1 | immature | A vs O | 12.027 | 0.000 |
| Afem.4100.1 | immature | A vs I | 12.027 | 0.000 |
| Afem.4100.1 | immature | I vs O | 0.000 | 1.000 |
| Afem.4100.1 | immature | A vs male | 3.076 | 0.195 |
| Afem.4100.1 | immature | I vs male | -8.951 | 0.003 |
| Afem.4100.1 | immature | O vs male | -8.951 | 0.003 |
| Afem.4100.1 | mature | A vs O | 13.910 | 0.000 |
| Afem.4100.1 | mature | A vs I | 13.910 | 0.000 |
| Afem.4100.1 | mature | I vs O | 0.000 | 1.000 |
| Afem.4100.1 | mature | A vs male | 3.397 | 0.158 |

*(continued)*

| Transcript | Stage | Contrast | log fold change | P-value |
|---|---|---|---|---|
| Afem.4100.1 | mature | I vs male | -10.512 | 0.001 |
| Afem.4100.1 | mature | O vs male | -10.512 | 0.001 |
| Afem.4103.1 | immature | A vs O | 10.897 | 0.001 |
| Afem.4103.1 | immature | A vs I | 5.110 | 0.049 |
| Afem.4103.1 | immature | I vs O | 5.787 | 0.025 |
| Afem.4103.1 | immature | A vs male | 1.311 | 0.554 |
| Afem.4103.1 | immature | I vs male | -3.798 | 0.119 |
| Afem.4103.1 | immature | O vs male | -9.586 | 0.002 |
| Afem.4103.1 | mature | A vs O | 13.596 | 0.000 |
| Afem.4103.1 | mature | A vs I | 10.063 | 0.002 |
| Afem.4103.1 | mature | I vs O | 3.533 | 0.132 |
| Afem.4103.1 | mature | A vs male | 3.292 | 0.167 |
| Afem.4103.1 | mature | I vs male | -6.771 | 0.016 |
| Afem.4103.1 | mature | O vs male | -10.304 | 0.002 |
| Afem.4111.2 | immature | A vs O | -0.624 | 0.660 |
| Afem.4111.2 | immature | A vs I | -0.299 | 0.833 |
| Afem.4111.2 | immature | I vs O | -0.326 | 0.818 |
| Afem.4111.2 | immature | A vs male | 0.782 | 0.584 |
| Afem.4111.2 | immature | I vs male | 1.081 | 0.451 |
| Afem.4111.2 | immature | O vs male | 1.406 | 0.330 |
| Afem.4111.2 | mature | A vs O | -0.258 | 0.856 |
| Afem.4111.2 | mature | A vs I | 0.409 | 0.773 |
| Afem.4111.2 | mature | I vs O | -0.667 | 0.639 |
| Afem.4111.2 | mature | A vs male | 1.659 | 0.255 |
| Afem.4111.2 | mature | I vs male | 1.250 | 0.387 |
| Afem.4111.2 | mature | O vs male | 1.916 | 0.192 |

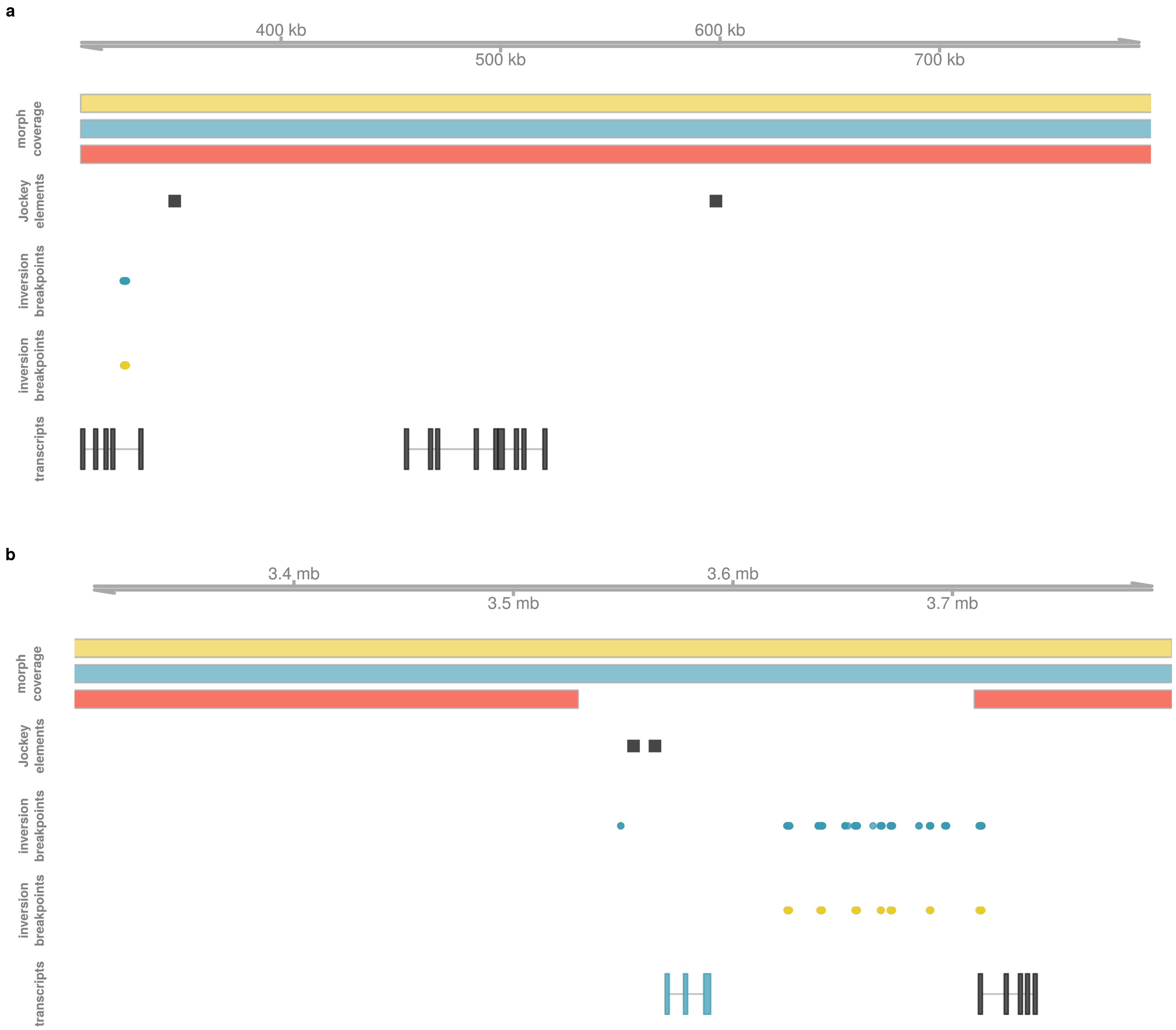

**Figure S12.** The regions that differentiate $I$ vs $O$ and $I$ vs $A$ females are situated $\sim$ 3.5 mb apart in the $I$ assembly (unlocalized scaffold 2 of chromosome 13) **a** In $I$-assembly coordinates, the region that differentiates $I$ vs $A$ females is located between 0.4 mb and 0.6 mb and is flanked by an uncharacterized gene (LOC124172704) at $\sim$ 0.3 mb. **b** A likely paralogue of this gene (LOC124172753) is located at $\sim$ 3.8 mb, flanking the presumed translocation of $A$ content. LINE annotations are located at $\sim$ 3.55 mb, on the other flank of the putative translocation. Each panel shows from top to bottom: morph-specific read depth coverage, the location of LINE retrotransposons in the the Jockey family, the mapping locations of $A$-derived and $I$-derived reads with a previously detected inversion signature against $O$ females, and transcripts expressed in $A$ and $I$ females of *I. elegans* and $A$ females of *I. senegalensis*. Black transcripts are present in both $A$ and $O$ haplotypes, and the blue transcript is uniquely present and expressed in $A$ and $I$.

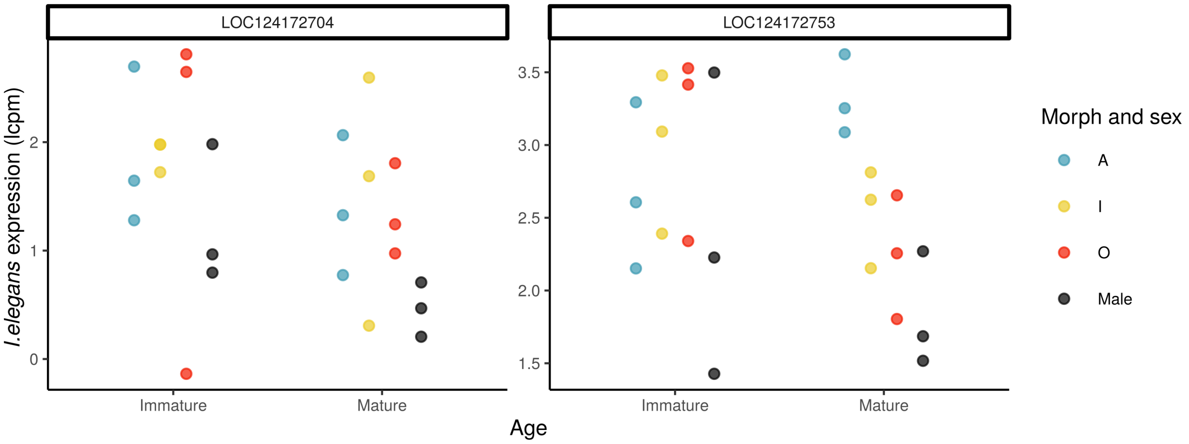

**Figure S13.** Expression of transcripts encoded by two similar and uncharacterized genes (LOC124172704 and LOC124172753) that border the morph locus in the *A* and *I* assemblies. Transcriptomic data were mapped to the DToL reference assembly, which includes the *O* haplotype and complete gene models. Expression was quantified as log counts per million and normalized across samples in adult females and males of *I. elegans*. P-values were computed from two-tailed exact tests and adjusted for multiple comparisons using the Benjamini and Hochberg's false discovery rate. There were no statistically significant differences in expression between groups for either gene (all FDR adjusted p-values $> 0.05$)

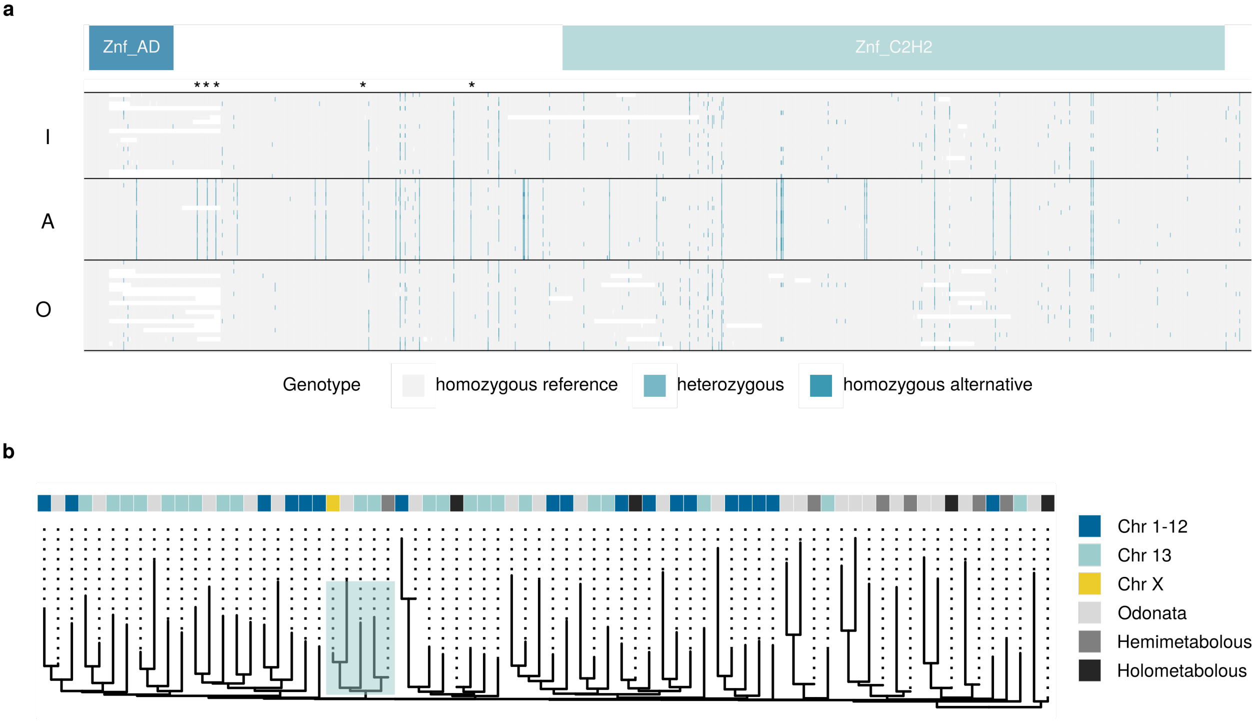

**Figure S14.** The candidate gene model Afem.4111 in *Ischnura elegans* **a** Genotype plot of the coding sequence of transcript Afem.4111.2. AD and C2H2 zinc-finger domain regions are highlighted in colour. The asterisks show non-synonymous mutations that distinguish *A* females from both *I* and *O* females. **b** Gene tree of Afem.4111 and its insect orthologues. The colour scale shows the chromosomal location of paralogues in *I. elegans* and the insect group in which othologues are found. "Odonata" refers to the Scarce chaser *Ladona fulva*, "Hemimetabolous" includes de proteomes of the Pea aphid (*Acyrthosiphon pisum*), the Western Flower thrip (*Frankliniella occidentalis*) and the Nevada termite (*Zootermopsis nevadensis*). "Holometabolous" includes the proteomes of the Squinting Brush Brown (*Bicyclus anynana*), the Flour beetle (*Tribolium castaneum*) and the Fruit fly (*Drosophila melanogaster*). The clade highlighted in blue includes both Afem.4111 transcripts.
